# Genome-scale metabolic model guided metabolic flux analysis in the endophyte *Alternaria burnsii* NCIM1409

**DOI:** 10.1101/2025.10.28.685040

**Authors:** Divya Dharshini Uma Shankar, Sarayu Murali, Shagun Shagun, Shyam Kumar Masakapalli, Karthik Raman, Smita Srivastava

## Abstract

Camptothecin (CPT), a potent anticancer alkaloid, is traditionally derived from plants like *Camptotheca acuminata* and *Nothapodytes nimmoniana*, but sustainable production remains challenging. This study explores the metabolic network of the fungal endophyte *Alternaria burnsii* NCIM1409, known for camptothecin production, using genome sequencing, genome scale metabolic modeling, and ^13^C-based pathway mapping. A genome-scale metabolic model (AltGEM *i*DD1552) was reconstructed, comprising 2188 reactions, 2148 metabolites, and 1552 genes, along with manual curation to include camptothecin biosynthesis pathways. Flux balance analysis identified key enzymatic targets, including secologanin synthase and tryptophan decarboxylase, for enhancing CPT production. Further, metabolic analysis of *A. burnsii* subjected to 20%[U-^13^C_6_] glucose or 99% [1-^13^C] glucose revealed active glycolysis, the pentose phosphate pathway, and the TCA cycle. This integrative approach provides insights into *A. burnsii*’s metabolic capabilities and highlights strategies for optimizing camptothecin biosynthesis, offering a foundation for sustainable production methods.

## Introduction

Camptothecin (CPT) is a pentacyclic monoterpene indole alkaloid predominantly found in two plants - *Camptotheca acuminata and Nothapodytes nimmoniana*^1^. It is one of the most potent indole alkaloids with anti-cancer properties, acting on DNA topoisomerase to induce cell death^2,3^. Camptothecin analogs, such as topotecan and irinotecan, have been successfully commercialized as anti-cancer drugs^3,4^.

Endophytes are microorganisms (often bacteria or fungi) that live inside different plant parts such as roots, stems, barks, leaves, and petiole^5,6^.These organisms cause no harm to the plant tissues and reside in plants as endosymbionts^7–9^. Endophytic fungi isolated from host plants have the potential to produce high-value bioactive compounds by the horizontal gene transfer (HGT) hypothesis^1^. Their ability to produce these bioactive metabolites has garnered significant interest all over the world.

Harvesting camptothecin from natural sources has significantly depleted the population of tree species in their native habitats. The synthetic production of camptothecin via chemical methods is tedious due to its intricate structure, characterized by multiple cyclic carbon rings. Furthermore, *in vitro* production from plant cell cultures is challenging, time-consuming and prone to microbial contamination. With increasing demand, endophytes offer a sustainable alternative for producing plant secondary metabolites. However, continuous subculturing of these endophytes under sterile conditions has led to attenuation in the production of secondary metabolites. Recently we reported that the fungal endophyte *Alternaria burnsii* NCIM1409, isolated from *N. nimmoniana*, has the potential to sustainably produce camptothecin under *in vitro* conditions^10^.

Given that *A. burnsii* can serve as a cell factory for camptothecin, defining its genome scale metabolic model including the key pathway nodes that regulate flux towards the camptothecin biosynthesis pathway will provide deeper insights into the organism’s biochemistry. This knowledge could lead to the development of strategies to further enhance camptothecin production.

Genome-scale metabolic model reconstruction (GSMM) involves a computational approach to map the complete set of metabolic reactions within an organism, based on its genome sequence. This method involves compiling a comprehensive list of genes, enzymes and metabolic pathways to create a network model that stimulates the organism’s metabolic network. GSMM has been effectively utilized to investigate the metabolic potential of various fungi and endophytes. For example, researchers have employed GSMM to study the specialized function of secondary replicons in host-associated niche adaptation of the legume symbiont *Sinorhizobium meliloti*^11^. Although no specific reports exist on the application of GSMM for camptothecin production in endophytes, it has been utilized for resveratrol production in *Alternaria* sp., particularly through the reconstruction of *Alternaria* sp. MG1^12^. These instances illustrate the usefulness of GSMM in understanding the metabolic capabilities of fungi and endophytes, which can guide metabolic engineering efforts and strain improvement.

In this study, we reconstructed GSMM of *Alternaria burnsii* NCIM1409 and combined it with metabolic flux balance analysis to elucidate the biosynthesis of camptothecin. Additionally, we incorporated ^13^C tracer experiments to gain a deeper understanding of the intracellular fluxes involved in the metabolism of *A. burnsii*. This integrated approach allowed us to map the metabolic pathways more accurately and identify potential strategies for enhancing CPT production. Adapted from the host-plant based on the HGT theory, a hypothetical CPT metabolism in *A. burnsii* NCIM 1409 (**Figure 1**).

**Figure 1:**
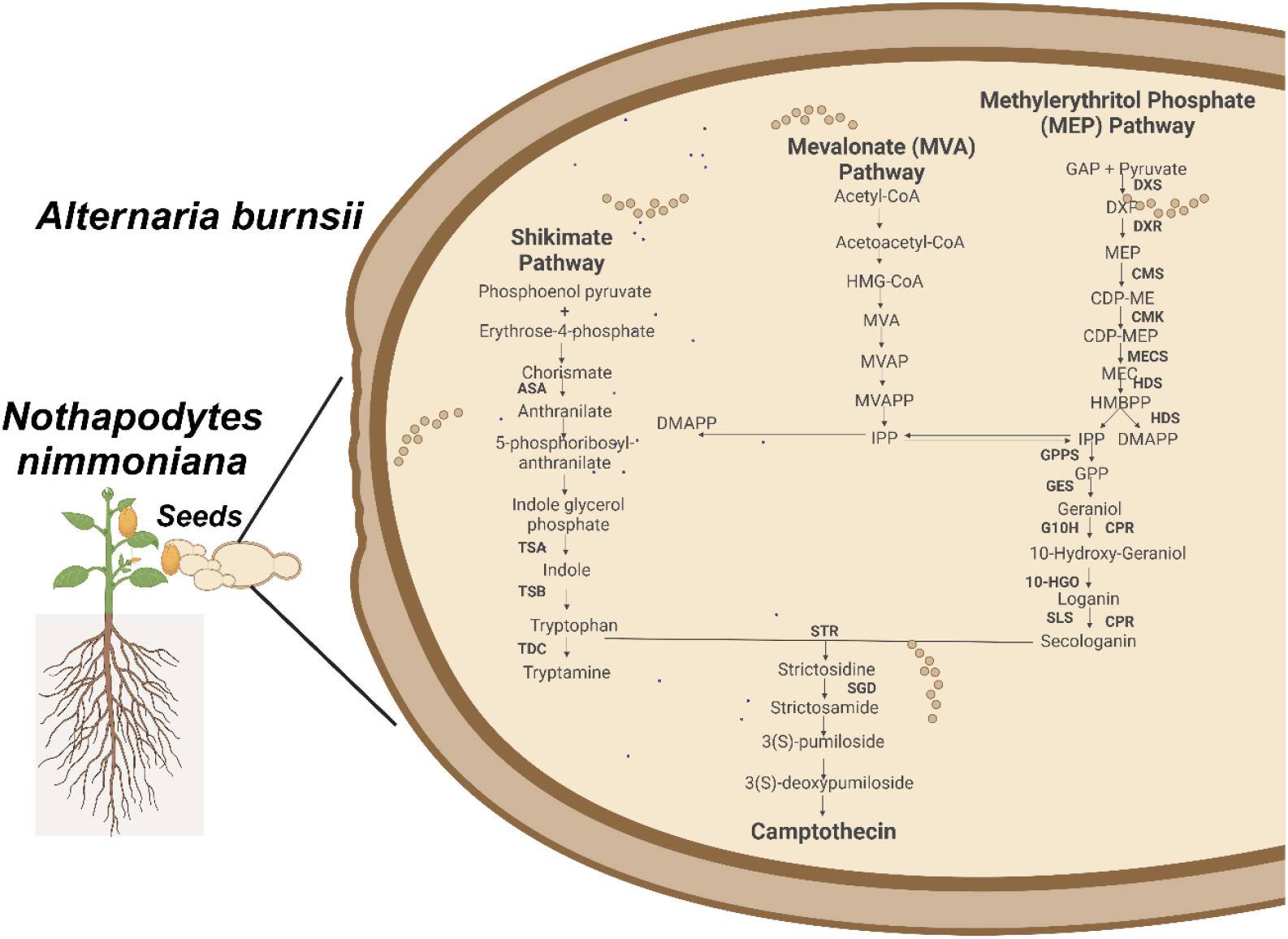
**Hypothetical Camptothecin pathway in *Alternaria burnsii* NCIM1409**, IA-Indole alkaloid, AS-Anthranilate synthase, TSB-Tryptophan synthase beta subunit, TDC-Tryptophan decarboxylase, DXR-deoxy-D-xylulose 5-phosphate reductoisomerase, HDR-Hydroxymethylbutenyl diphosphate reductase, IPI-Isopentenyl diphosphate isomerase, GPPS-Geranyl diphosphate synthase, GES-Geraniol synthase,G10H-Geraniol 10-hydroxylase,10 HGO10-hydroxygeraniol oxidoreductase, IS-Iridoid synthase, SLS-Secologanin synthase, CPR NADPH-cytochrome P450 reductase, STR-Strictosidine synthase, SGD-Strictosidine beta-D-glucosidase.

## Results

### Genome sequence and features reveals characteristics of *A. burnsii* NCIM1409

Previously sequenced, the genome of this *A. burnsii* NCIM1409 strain has a GC content of 43%, determined using a paired-end library^35^. The genome was characterized by several functional gene sequences (**Table 1**), including 78 genes prominently involved in secondary metabolite biosynthesis. The distribution of genes involved in metabolism, genetic information processing, and cellular processes is illustrated in **Figure 2A–C**

**Table 1:** Features of *A. burnsii* NCIM1409 genome sequence and properties of gene annotation (Eurofins Scientific, India). The table summarizes key metrics such as genome size, GC content, and gene annotation statistics, including Gene Ontology classifications.

| GENERAL FEATURES |  |
| --- | --- |
| Genome size (Mb) | 3,32,35,878 |
| GC content (%) | 43.35 |
| Scaffold N50 (bp) | 8,32,062 |
| Number of protein-coding sequences | 13,606 |
| ORF coverage (%) | 12 |
| PROPERTIES OF GENE ANNOTATION |  |
| Total Genes | 10,504 |
| Genes with Gene Ontology | 7,071 |
| Genes with Biological processes | 2,491 |
| Genes with Cellular Components | 1,641 |
| Genes with Molecular Function | 2,939 |

**Figure 2:**
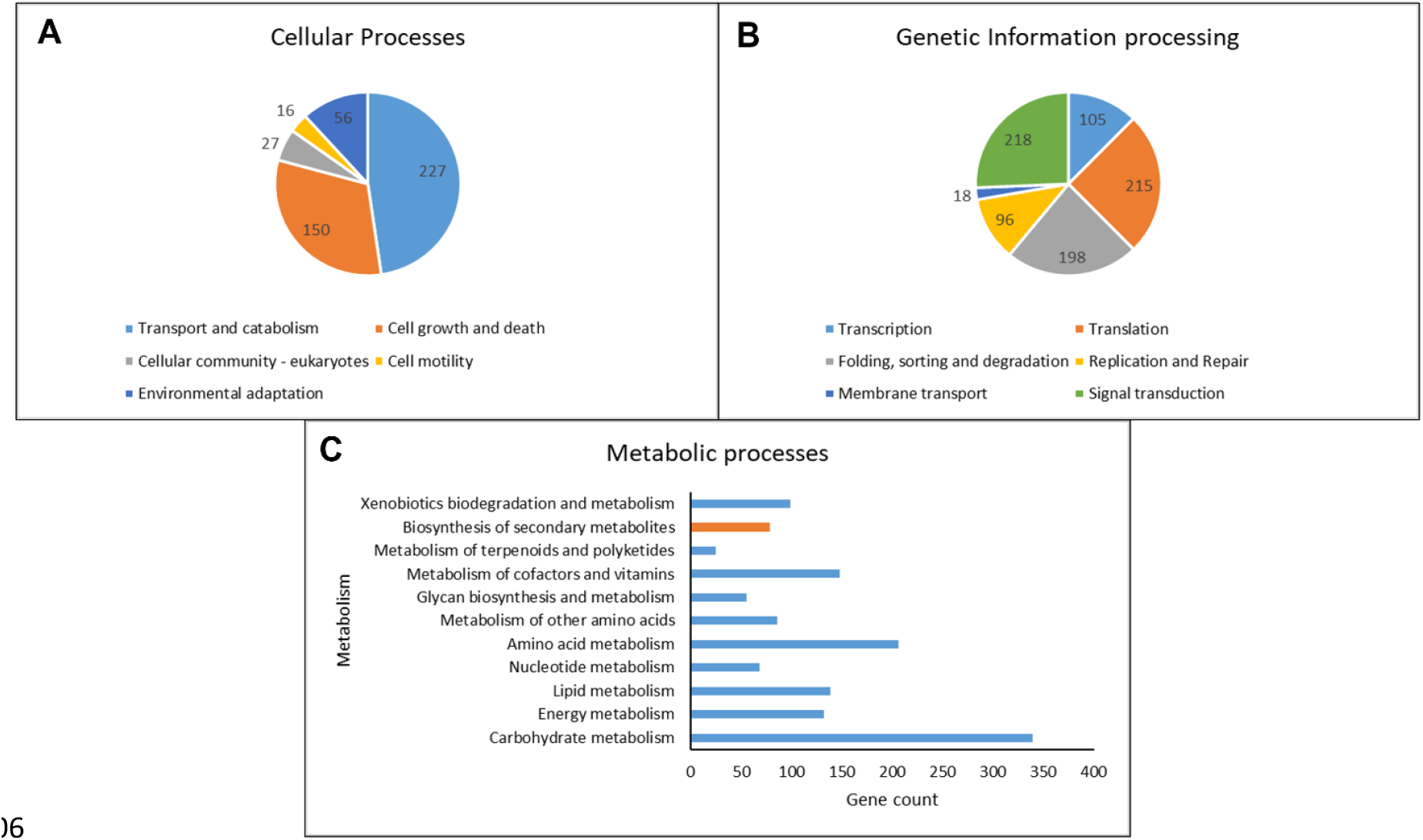
Genes involved in metabolism, genetic information processing, and cellular processes. The different genes in **A**. Cellular Processes: Transport and catabolism (227 genes), cell growth and death (150 genes), cellular community in eukaryotes (27 genes), cell motility (16 genes), and environmental adaptation (56 genes); **B**. Genetic Information Processing: Transcription (105 genes), translation (215 genes), folding, sorting, and degradation (198 genes), replication and repair (96 genes), membrane transport (18 genes), and signal transduction (218 genes) and **C**. Metabolic Pathways: Metabolic processes, including carbohydrate metabolism (highest gene count), energy metabolism, lipid metabolism, nucleotide metabolism, amino acid metabolism, and others.

### A Curated Genome-Scale Model Captures Core and Secondary Metabolism

In the initial construction of the metabolic model for *A. burnsii*, the ModelSEED platform was utilized to generate an automated draft. This preliminary model necessitated substantial manual curation to ensure accuracy and completeness. The availability of the whole-genome sequence and corresponding sequence annotation for *A. burnsii* facilitated this curation process^35^(Eurofins Scientific, India). The draft model initially contained 906 gene-protein associations formatted according to the PATRIC database^36^. These associations were subsequently converted to the NCBI format, which resulted in 640 reactions remaining without any gene-protein-reaction associations. The draft model initially employed the SEED format, which included some degenerate metabolites. To address this issue, the model was converted to the BiGG Models^20^ namespace. This conversion enabled easier reaction mapping in Escher and provided metabolite identifiers that are more easily recognizable and follow a standardized nomenclature^37^. A significant challenge in the manual reconstruction process was the operability of gap-filling. The draft model contained 953 gaps with limited pathway connections. The manual reconstruction of pathways involved in secondary metabolism further increased the number of gaps. Therefore, the gap-filling framework was implemented manually from top to bottom, ensuring the correct directionality of metabolic reactions.

Following extensive manual refinements, a comprehensive GSMM for *Alternaria burnsii* was constructed. This model termed – *Alt*GEM *i*DD1552, includes 2,188 reactions, 2,148 metabolites, and 1,552 genes, representing 11.31% of the total protein-coding genes (**Figure 3 A, B)**. The model spans eight compartments: cytoplasm, vacuole, nucleus, extracellular space, peroxisome, mitochondria, endoplasmic reticulum, and other uncompartmentalized regions. The camptothecin biosynthetic pathway, which involves enzymes from the mevalonate EMP pathway, was localized to the cytoplasm. This localization suggests that camptothecin biosynthesis occurs in the cytoplasm, and the compound is subsequently secreted into the extracellular space. The model is provided in the community standard SMBL (Level 3 Version 2 with FBC) and excel format in the supplementary material S1.

**Figure 3:**
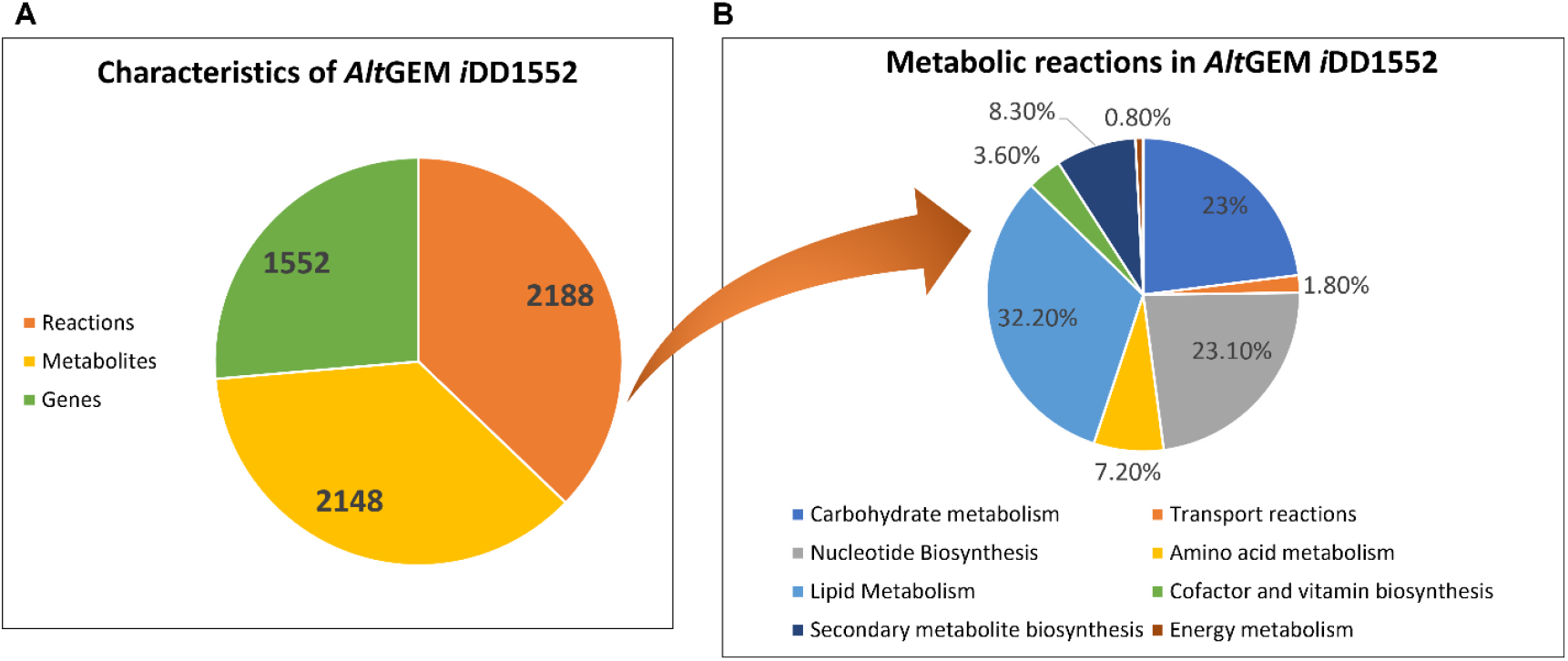
Characteristics of *Alt*GEM *i*DD1552. **A**. Overall distribution of homologous genes across three major taxonomic groups and **B**. Breakdown of one of the groups, highlighting the relative abundance of genes among different species.

The reconstructed metabolic model includes a biomass reaction, represented by functions of metabolites, energy molecules, nucleotides, lipids, and cell wall components. This reaction incorporates precursor molecules that lead to the formation of higher biomolecules, including proteins and long-chain carbohydrates. The biomass composition of each component in the equation was meticulously calculated, and stoichiometric coefficients were determined and fixed, as mentioned in the supplementary table 11. The biomass composition for lipids and mannitol was adapted from *Alternaria* sp. MG1^12^ and included in the supplementary material S1. As *Alternaria burnsii* NCIM1409 is a novel organism with no available experimental data, growth-associated maintenance (GAM) and non-growth-associated maintenance (NGAM) parameters were adapted from *Alternaria* sp. MG1^12^, amounting to 71.4 mmol ATP gDW^-1^ h^-1^.

Following the central carbon metabolism, the pathways leading to camptothecin were included and checked for *in-silico* flux along the pathway (**Figure 4**). The *in vitro* studies of *Alternaria burnsii* show that Potato Dextrose Broth (PDB) was used as the culture medium for *Alternaria burnsii*, unlike the other media, showing the less futile growth. The medium rich in proteins, carbohydrates, and other minerals were accounted and constrained in the model. The maximum substrate utilization rate and essential amino acids were constrained as -1.62 and -0.01 mmol gDW^-1^h^-1^, respectively in the model for flux balance analysis.

**Figure 4:**
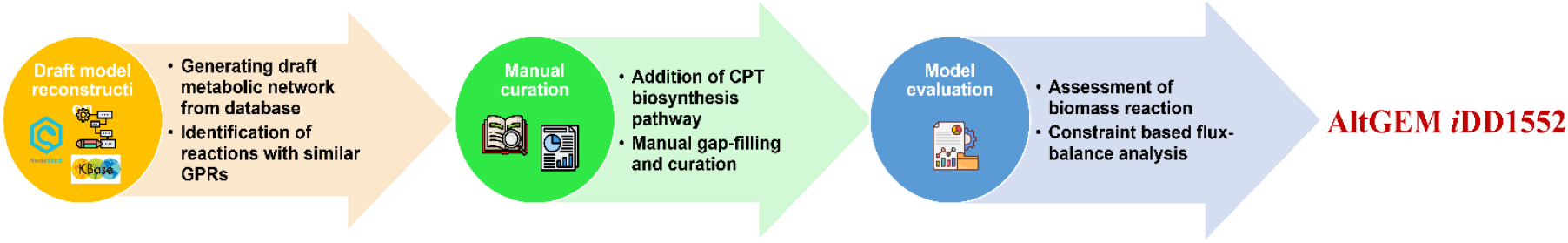
Workflow for metabolic model reconstruction. The process consists of three main stages: (i) Draft model reconstruction, where a draft metabolic network is generated from a database, and reactions with similar gene-protein-reaction (GPR) associations are identified; (ii) Manual curation, which includes the addition of the CPT biosynthesis pathway and manual gap-filling; and (iii) Model evaluation, involving the assessment of the biomass reaction and constraint-based flux balance analysis. The final curated model is designated as AltGEM *i*DD1552.

### Model Refinement Improves Predictive Accuracy and Validates Growth Estimates

To validate the model, it was constrained based on the experiments conducted on the organisms. All the media components and growth rates were converted to the standard format and were taken as the input to the model. Experimentally determined values for different substrates as shown in the supplementary Table 12 were tested to improve the model predictability. The growth of *Alternaria burnsii* was verified and compared using the experimental studies and the available data. Some of the model discrepancies include the ability of the strain to produce important components in biomass reaction, but it was not present in the gene annotation due to missing reactions, gaps, and other unknown pathways. The exchange reactions for Na^+^, K^+^, and Fe^2+^ were constrained since these are important for the growth of the organism. With these constrained values, the model predicted specific growth rate as 0.0288 h^−1^, which was consistent with the *in vitro* study value of 0.0233 h^−1^, with a 19.09% deviation. The metabolic characteristics and fermentation studies predict the capability of camptothecin production in this endophyte. Thus, the model was constrained using the eight different sources to increase the accuracy of the predicted values.

### Flux Analysis Identifies Enzymatic Targets for Enhanced Camptothecin Biosynthesis Secologanin Synthase and Strictosidine Synthase Drive Key Flux Toward Camptothecin

The FSEOF strategy was applied to the metabolic model of *Alternaria burnsii* to identify fluxes that increased with enforced camptothecin production, resulting in the prediction of 67 enzyme targets in the overall metabolic model. These targets were ranked and scored using the *f*_*PH*_ metric based on their strategic roles in directing flux toward camptothecin biosynthesis in the secondary metabolic pathway.

As seen in **Table 2**, secologanin synthase has a higher *f*_*PH*_score for overexpression enzyme targets. Secologanin synthase catalyzes the synthesis of strictosidine along with the tryptamine from the shikimate pathway present in the downstream pathway leading to camptothecin production. The overexpression studies with the enzyme secologanin synthase from *N. nimmoniana* showed a 2.02–2.86-fold increase in CPT accumulation ^38^. Other predicted targets are strictosidine synthase (the final step toward camptothecin production) and tryptophan decarboxylase (synthesis of tryptamine from tryptophan). The studies on overexpressing two important key genes in the Terpenoid indole pathway: Tryptophan decarboxylase and strictosidine synthase in *Catharanthus roseus* show an increase in antineoplastic drug metabolite vinblastine^39^. The downstream camptothecin biosynthetic pathway starts from the isoprenoid subunits and its intermediate geranyl diphosphate (GPP). In subsequent steps, it forms secologanin by hydroxylation, reduction, and glycosylation reactions that act as the key compound to synthesize various classes of other secondary metabolites.

**Table 2:** Enzymes showing significant overexpression in the Camptothecin biosynthesis. The table highlights ten reactions with high *f*_*PH*_, indicating their possible regulatory roles for the increase in the camptothecin production.

| Reaction Name | Metabolic Pathway | $f_{PH}$ |
| --- | --- | --- |
| Secologanin synthase | Camptothecin biosynthesis | 0.99 |
| Strictosidine synthase | Camptothecin biosynthesis | 0.97 |
| L-Tryptophan decarboxy-lyase | Shikimate Pathway | 0.93 |
| Tryptophan synthase (indoleglycerol phosphate) | Shikimate Pathway | 0.92 |
| NAD<sup>+</sup> kinase | Camptothecin biosynthesis | 0.85 |
| 2-C-methyl-D-erythritol 4-phosphate cytidyltransferase | Camptothecin biosynthesis | 0.84 |
| 1-Deoxy-D-xylulose-5-phosphate reductoisomerase | Camptothecin biosynthesis | 0.83 |
| (S)-8-Oxocitronellyl enol synthase | Camptothecin biosynthesis | 0.82 |
| 8-Hydroxygeraniol dehydrogenase | Camptothecin biosynthesis | 0.81 |

### Redirecting Central Carbon Metabolism Enhances Camptothecin Yield

These predicted targets were ranked based on the *f*_*PH*_ score, to maintain biomass and camptothecin production balance. Table 3 shows that the aldehyde dehydrogenase has a higher *f*_*PH*_ score for knockout studies. Aldehyde dehydrogenase (ADH2) catalyzes the synthesis of ethanol from acetaldehyde^40^.

**Table 3:** Enzymes showing significant knockout targets in the Camptothecin biosynthesis. The table highlights ten reactions with high *fPH*, indicating their possible regulatory knockout roles for the increase in the camptothecin production.

| Enzyme name | Metabolic Pathway | $f_{PH}$ |
| --- | --- | --- |
| Aldehyde dehydrogenase | Central carbon metabolism | 0.99 |
| Formaldehyde:NAD<sup>+</sup> oxidoreductase | Central carbon metabolism | 0.99 |
| Formaldehyde dehydrogenase | Central carbon metabolism | 0.97 |
| D-xylulose 5-phosphate D- glyceraldehyde-3-phosphate-lyase (adding phosphate; acetyl-phosphate- forming) | Pentose phosphate pathway | 0.92 |
| Phosphoserine aminotransferase | Central carbon metabolism | 0.92 |
| Phosphoglycerate dehydrogenase | Central carbon metabolism | 0.90 |
| Acetate exchange | Central carbon metabolism | 0.89 |
| Acetate kinase | Central carbon metabolism | 0.88 |
| Phosphotransacetylase | Central carbon metabolism | 0.88 |
| Acetate plasma membrane transport | Central carbon metabolism | 0.88 |

Deletion of ADH genes, particularly ADH1, reduces ethanol formation, a major carbon sink in yeast, and redirects pyruvate-derived carbon into the pyruvate dehydrogenase (PDH) bypass, which boosts cytosolic acetyl-CoA availability^41^. Other knockout targets, such as Acetate kinase and Formaldehyde dehydrogenase and those relating to the isoprenoid subunit pathway in fungi merits future investigations.

### ^13^C Labeling Confirms Active Glycolysis, PPP, and TCA Pathways Supporting Camptothecin Biosynthesis

The average ^13^C abundance and mass isotopomer distributions (MIDs) of various fragments of amino acids highlighted the activities of central metabolic pathways in *Alternaria burnsii* **(Figure 6 &7, Supplementary table 13 & 14)**. The average ^13^C in the proteinogenic amino acids of cells grown in media supplemented with 20% [U-^13^C_6_] glucose were low (1-3%) showing the higher contributions of other carbon sources from potato infusion media or pre-existing biomass **(Figure 6, Supplementary table 13)**. Specifically, the maximum average ^13^C was observed in Alanine [m/z 260 (3%); m/z 232 (4%)] while other amino acids showed very low ^13^C incorporation but higher than the unlabeled experiments. The average label incorporation in alanine [m/z 260 (3%)] and serine [m/z 390 (1%)] were higher than the natural abundance and hence confirm the metabolic activity of upper glycolysis and/or Pentose Phosphate Pathway (PPP). ^13^C incorporations in histidine [m/z 440 (1%)], aspartic acid [m/z 418 (2%)] and glutamic acid [m/z 432 (2%)] were low but seemed higher than natural abundance, thereby needing to examine the incorporations from [1-^13^C] glucose feeding experiments to make any reliable assessment of pathway activities.

**Figure 5:**
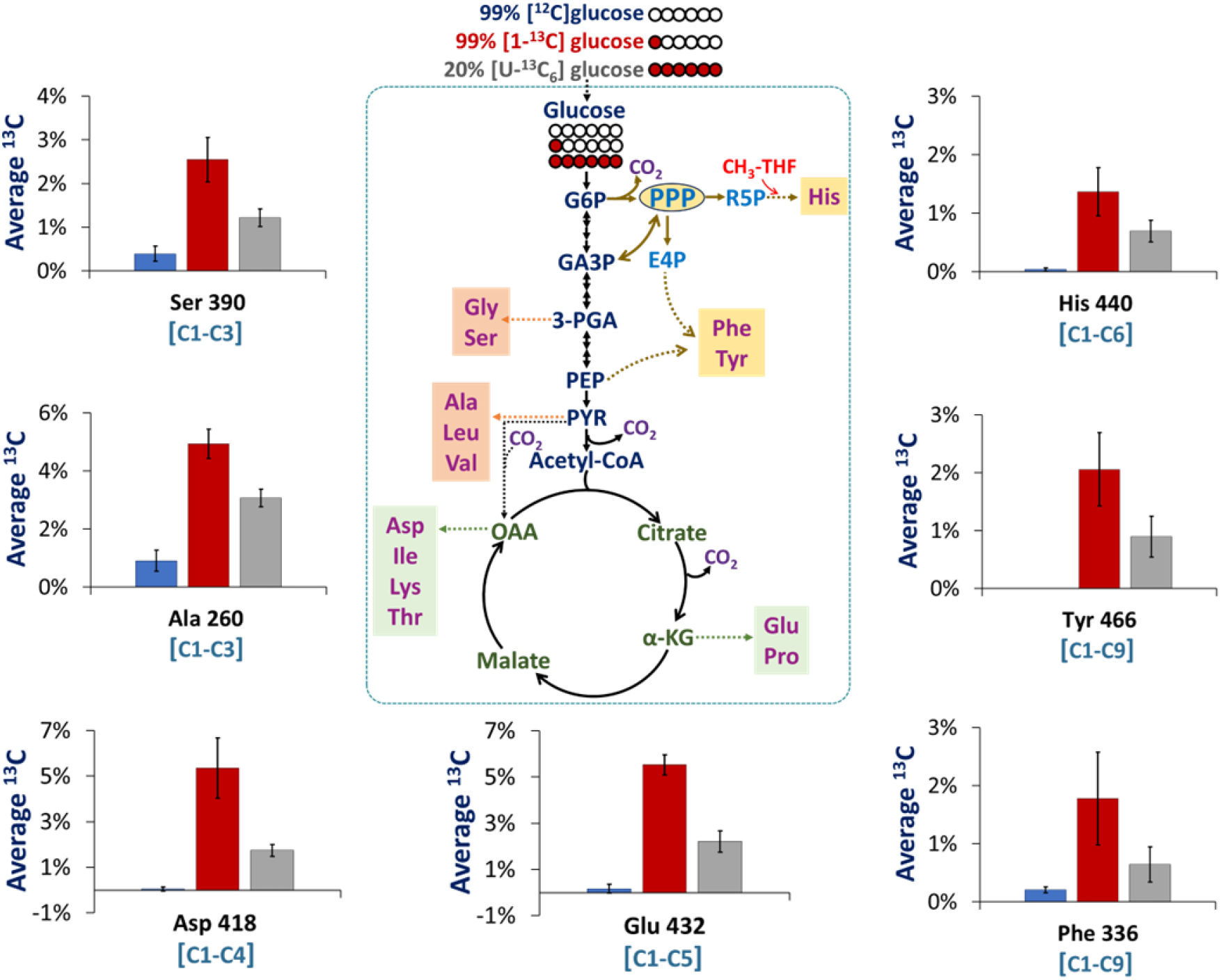
The average ^13^C incorporation in the valid amino acid fragments of *Alternaria burnsii* highlighted the activities of various central metabolic pathways present. In parallel experiments, A burnsii cells were grown in potato infusion media supplemented with either 99% [^12^C]glucose; 20% [U-^13^C_6_] glucose, or 99 % [1-^13^C]glucose. Average 13C incorporation in each metabolite fragment is presented in % age, represented as mean ± standard deviation (SD); n = 4] [Abbreviations used are G6P: Glucose-6-Phosphate; GA3P: Glyceraldehyde 3-Phosphate; 3-PGA: 3-Phosphoglyceric acid; PEP: Phosphoenolpyruvate; PYR: Pyruvate; PPP: Pentose Phosphate Pathway; R5P: Ribose-5-Phosphate; E4P: Erythrose-4-Phosphate; OAA-Oxaloacetic acid; α-KG: alpha-Ketoglutarate; CH_3_-THF: Methyl-Tetrahydrofolate; Ser: Serine; Ala: Alanine; His: Histidine; Tyr: Tyrosine; Asp: Aspartic acid; Glu: Glutamic acid; Phe: Phenylalanine].

**Figure 6:**
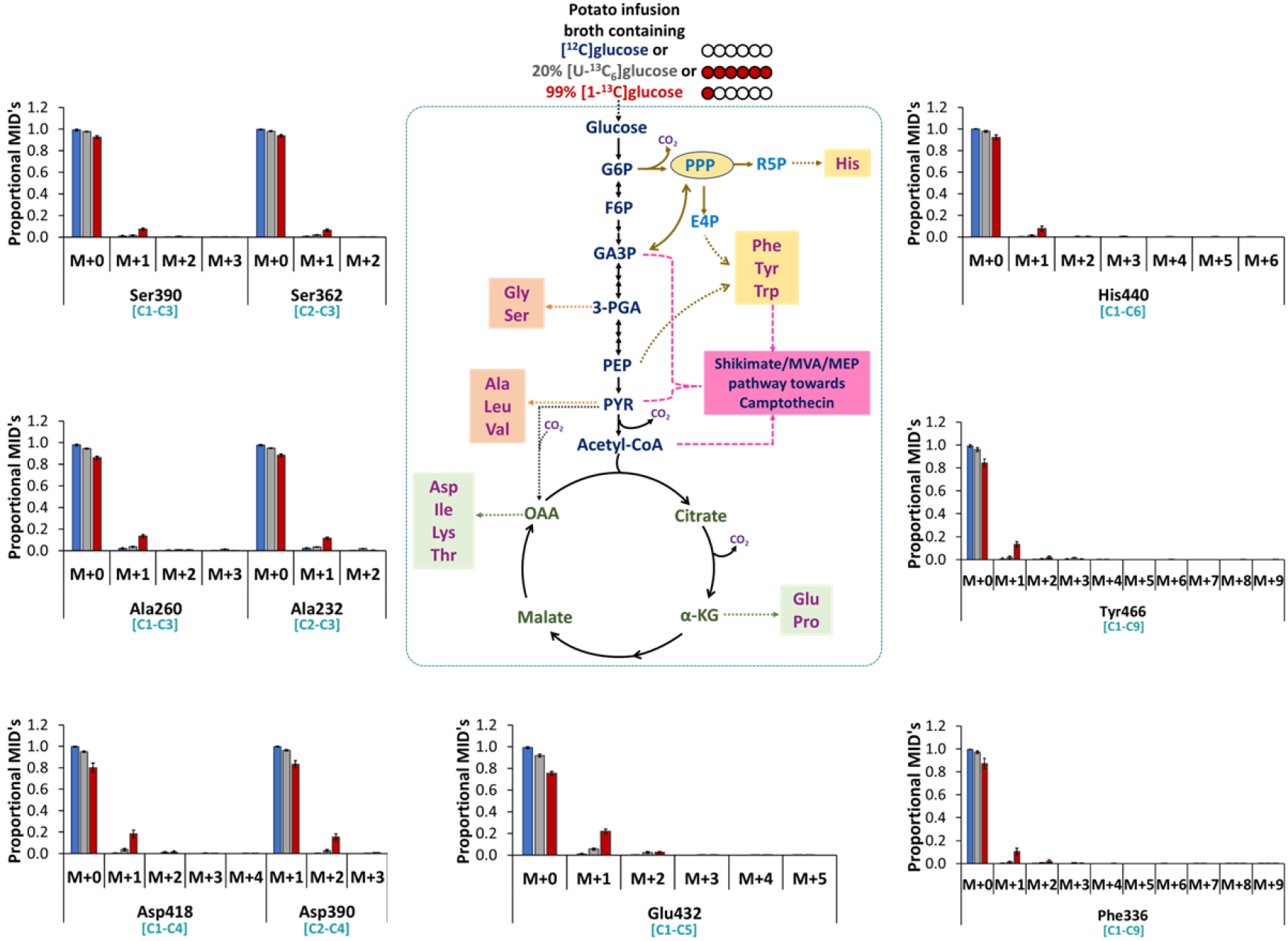
The mass isotopomer distributions (MIDs) of the valid amino acid fragments of of *Alternaria burnsii* highlighted the activities of various central metabolic pathways. The cells were grown in potato infusion media supplemented with 99% [^12^C]glucose; 20% [U-^13^C_6_]glucose, and 99 % [1-^13^C]glucose. The proportional MIDs of each metabolite fragment is represented as mean ± standard deviation (SD); n = 4 [Abbreviations used are G6P: Glucose-6-Phosphate; GA3P: Glyceraldehyde 3-Phosphate; 3-PGA: 3-Phosphoglyceric acid; PEP: Phosphoenolpyruvate; PYR: Pyruvate; PPP: Pentose Phosphate Pathway; R5P: Ribose-5-Phosphate; E4P: Erythrose-4-Phosphate; OAA-Oxaloacetic acid; α-KG: alpha-Ketoglutarate; CH_3_-THF: Methyl-Tetrahydrofolate; Gly: Glycine; Ser: Serine; Ala: Alanine; Leu: Leucine; Val: Valine; Asp: Aspartic acid; Ile: Isoleucine; Lys: Lysine; Thr: Threonine; Glu: Glutamic acid; Pro: Proline; Phe: Phenylalanine; Tyr: Tyrosine; Trp: Tryptophan; His: Histidine; MVA: Mevalonate; MEP: Methylerythritol phosphate].

Cells grown in 99% [1-^13^C] glucose, showed average ^13^C of 1-6% incorporation in the proteinogenic amino acids. The label incorporation in alanine [m/z 260 (5%)], serine [m/z 390 (3%)], histidine [m/z 440 (1%)], aspartic acid [m/z 418 (5%)] and glutamic acid [m/z 432 (6%)] confirmed active glucose oxidation via the central metabolic pathways. The low levels of ^13^C in histidine can be attributed to the loss of C1 from glucose via the PPP. Further, the average ^13^C incorporation in aromatic amino acids – Phenylalanine and Tyrosine (2%) highlight the contribution of carbon from glucose towards shikimate pathway in *A. burnsii* which could be of interest. Over, the ^13^C metabolic analysis confirmed the pathway activities of glycolysis, PPP and TCA cycle **(Figure 5, Supplementary table 13)**. These findings were further supported by the MIDs of the amino acid fragments from the positional feeding experiments **(Figure 6, Supplementary table 14)**.

## Discussion

The genome sequencing of *Alternaria burnsii* offers a detailed understanding of its metabolic and functional capabilities, showcasing its potential for biotechnological applications, particularly in secondary metabolite biosynthesis. With a GC content of 43.35%, the genome aligns with other *Alternaria* species, indicating a balance between genomic stability and adaptability. The prediction of 12,000 genes highlights the organism’s complex biosynthetic repertoire, further underscored by the identification of 78 genes dedicated to secondary metabolite production. This suggests a significant potential for producing bioactive compounds, notably camptothecin, a valuable anti-cancer alkaloid (**Figure 1**).

The functional annotation of *A. burnsii* revealed 7,071 genes with Gene Ontology (GO) terms, categorizing them into biological processes, cellular components, and molecular functions. This annotation aids in predicting metabolic pathways and provides insights into the organism’s biosynthetic potential (**Figure 2**). The high number of genes linked to secondary metabolism underscores *A. burnsii*’s potential as a source of pharmaceutical compounds. The presence of genes associated with the biosynthesis of camptothecin highlights its relevance for metabolic engineering.

The genome-scale metabolic model (AltGEM iDD1552) developed for *Alternaria burnsii* NCIM1409 provides a systems-level framework to understand the organism’s metabolic potential and its role in camptothecin biosynthesis (**Figure 3**). The reconstruction integrates genomic, biochemical, and physiological data to predict feasible flux distributions and growth behavior under defined conditions. Model refinement and validation against experimental data improved its predictive accuracy, establishing confidence in its use for metabolic exploration (**Figure 4**). Constraint-based simulations, including flux balance analysis and FSEOF, identified key enzymes whose modulation could enhance camptothecin production, linking primary metabolism with secondary metabolite pathways. The model further enables hypothesis generation for precursor supply, cofactor balancing, and alternative nutrient utilization.

The genome-scale metabolic model identified several enzymes as pivotal targets for enhancing camptothecin biosynthesis (**Table 2, 3**). Among these, secologanin synthase emerged as a critical enzyme, catalyzing the synthesis of strictosidine—a precursor to camptothecin. Studies on *N. nimmoniana* have shown that overexpression of secologanin synthase can significantly increase camptothecin accumulation, suggesting that similar strategies could be applied to *A. burnsii*. Other targets, such as strictosidine synthase and tryptophan decarboxylase, play essential roles in the biosynthetic pathway and have demonstrated success in enhancing secondary metabolite production in other systems. The upstream isoprenoid pathway, responsible for producing secologanin via geranyl diphosphate (GPP), also represents a crucial area for intervention. Enzymes such as NAD+ kinase and DXR are vital for this pathway, with previous studies demonstrating their efficacy in boosting secondary metabolite yields through metabolic engineering. Knocking out key metabolic genes such as alcohol dehydrogenase (ADH), acetate kinase, and formaldehyde dehydrogenase (SFA1) in *Saccharomyces cerevisiae* has been shown to enhance carbon flux toward acetyl-CoA, thereby increasing the biosynthesis of isoprenoid subunits. Deletion of ADH genes, particularly ADH1, reduces ethanol formation, a major carbon sink in yeast, and redirects pyruvate-derived carbon into the pyruvate dehydrogenase (PDH) bypass, which boosts cytosolic acetyl-CoA availability. This strategy has been experimentally validated to enhance the production of acetyl-CoA-derived compounds such as n-butanol, fatty acids, and various isoprenoids, including amorphadiene and santalene^41–44^

The use of ^13^C isotope labeling provided a detailed understanding of *A. burnsii*’s metabolic pathways. Labeling studies revealed active glycolysis, the pentose phosphate pathway (PPP), and the TCA cycle, with alanine,serine, histidine, aspartic acid and glutamic acid [m/z 432 (6%)] label incorporation confirming these activities (**Figure 5, 6**). The involvement of the shikimate pathway in aromatic amino acid biosynthesis further highlights its significance in camptothecin production. Similar findings in fungi like *Trichoderma reesei* ^45^ and *Ashbya gossypii* ^46^ corroborate these results, emphasizing the utility of isotope-labeling studies in elucidating metabolic fluxes.

This study establishes a robust platform for optimizing camptothecin biosynthesis in *A. burnsii*. By integrating genome sequencing, metabolic modeling, and experimental data, it identifies actionable enzyme targets and proposes strategies for metabolic reprogramming. Future work should focus on validating these targets through gene overexpression and knockout studies, potentially leveraging CRISPR-Cas9 for precise genome editing. Such advancements could address the ecological and industrial challenges associated with camptothecin production, paving the way for sustainable production methods. A genome-scale metabolic model was reconstructed for camptothecin-producing endophytic fungus, *Alternaria burnsii*. The verification of the occurrence of the camptothecin pathway was investigated in a non-genetically modified microorganism at the gene level. The diverse capabilities and possible key genes in *Alternaria burnsii* to produce camptothecin were studied. Furthermore, the key genes involved in the improvement of camptothecin production extend towards metabolic intervention studies. Systems biology has become an essential and useful strategy for the creation of platform strains and help in the microbial bioproduction of secondary metabolites. The reconstruction of genome-scale metabolic models has facilitated the reckoning of the mechanistic behavior that exists, which is difficult to culture and predict its growth. The major challenge would be the mechanism involved in the non-native organism. A computational-guided approach for understanding *Alternaria burnsii* is a powerful tool to garner metabolic pathways and interconnections. The combinatorial effort of using ^13^C Metabolic Flux Analysis (MFA) to quantify intracellular fluxes step closer to the comparison of predicted values in the model and combats the challenge. Based on the analysis, it can be further extended toward the rational metabolic engineering studies to tweak the camptothecin production.

## Limitations of the study

While the genome-scale metabolic model (AltGEM *i*DD1552) of *Alternaria burnsii* NCIM1409 provides valuable insights into its metabolic capabilities and potential routes for camptothecin biosynthesis, certain limitations remain. The model’s predictive power is constrained by the completeness and accuracy of current genome annotations and the limited availability of organism-specific kinetic and regulatory data. Experimental validation of all target enzymes was not feasible within the scope of this study. Moreover, the model assumes steady-state conditions and does not capture dynamic regulatory effects or compartmentalized metabolite transport in detail. Future integration of transcriptomic, proteomic, and time-resolved flux data will be essential to enhance the model’s predictive accuracy and biological relevance.

## Star★Methods

### Genome assembly and annotation

For the reconstruction of the GSMMs, paired-end sequencing was performed on the metagenomic DNA sample using Illumina TruSeq Nano DNA Library Prep Kit. The Augustus (version.3.3.3) gene prediction tool was used to determine the assembly of the genome, functional analysis of genes, and genome annotation. Predicted gene sequences were searched against the NCBI non-redundant protein database (nr) using the Basic Local Alignment Search Tool (BlastX) with an E-value threshold of 1e-05, facilitated by the Diamond tool. This process annotated 9,879 genes, while 625 genes had no significant BLAST hits. Most of the BLAST hits corresponded to homologs in *Alternaria alternata*. A summary of the BLAST results is presented in **Figure 7**.

**Figure 7:**
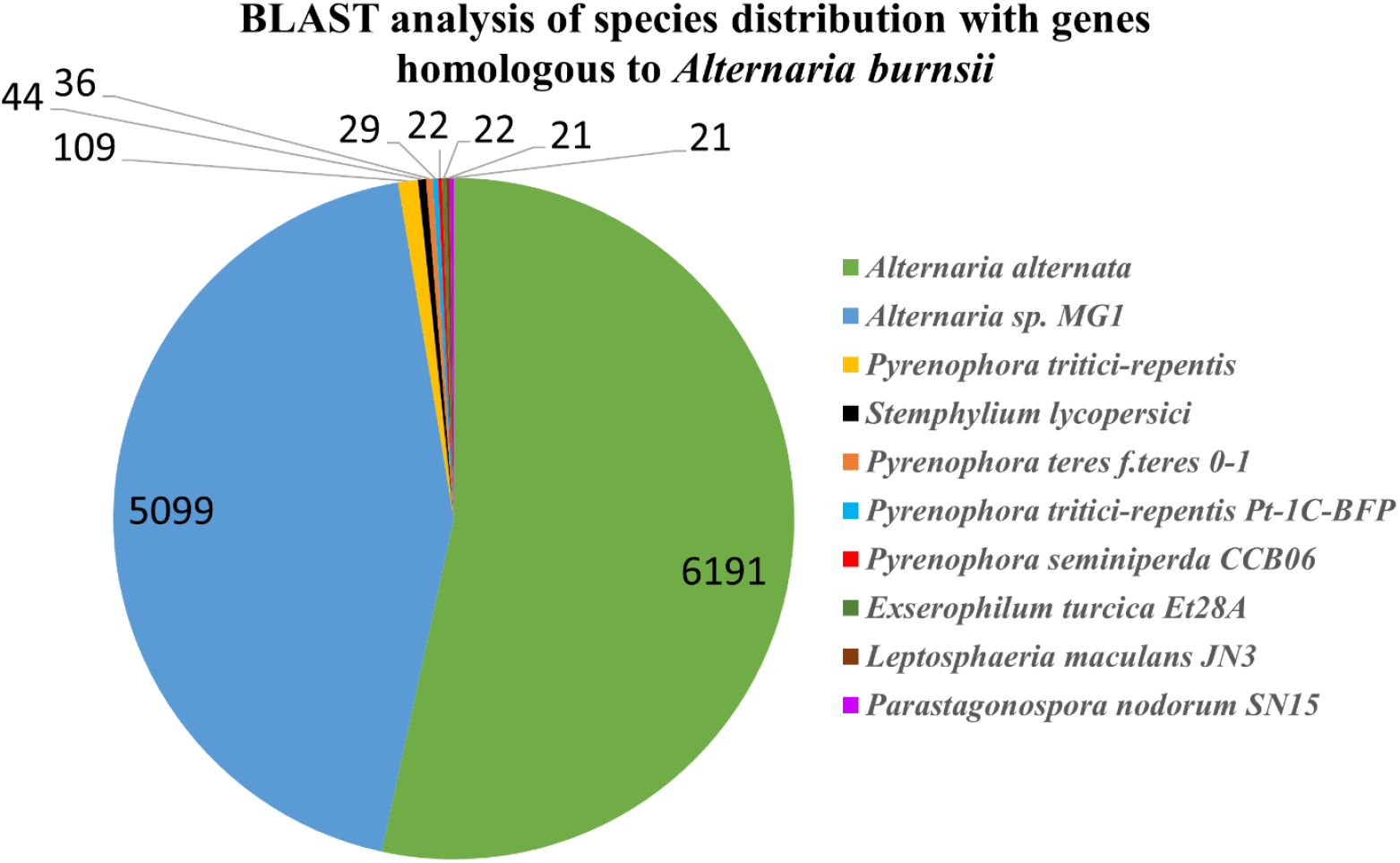
BLAST analysis showing the species distribution of genes homologous to *Alternaria burnsii* NCIM1409. Different species are represented in the pie chart, indicating the relative proportion of homologous genes found in each.

### Metabolic reconstruction

The objective of this study was to develop a metabolic model to interpret the metabolism of *A. burnsii*. To develop a draft reconstruction, genomic information and annotated sequences of *A. burnsii* were collected from different biochemical databases like KEGG^13^ and Biocyc^14^. The KEGG database was employed to confirm the specific metabolic capabilities and assemble unique reactions that were absent in reference strains. The MetaCyc^15^ database was also utilized to verify the reaction reversibility. Additionally, the TCDB^16^ (Transporter Classification Database) was consulted to obtain transport reactions. Automated reconstruction was facilitated through the ModelSEED^17^ database and simulated using the COBRA toolbox 3.0^18^ and SBML toolbox incorporated in MATLAB (version 2024a, MathWorks Inc.).

The draft model generated by ModelSEED comprised a biomass reaction which accounted for protein, DNA, and RNA. However, specific biosynthesis reactions leading to cofactor components and growth maintenance reactions were added to fill the missing information and metabolic connections in the network. The secondary metabolism leading to the production of camptothecin was added to the reconstruction from Plant metabolic networks database^19^. Corresponding gene-protein reactions, EC numbers and reaction sub-localization were cross-referenced from KEGG and BIGG databases. Following the mapping of metabolic reactions with their functions, the ModelSEED namespace was converted to the BIGG database namespace. This conversion was done to ensure consistency across the model, allowing for reproducibility and effective comparison with other existing fungal models. Further, manual curation involved ensuring stoichiometric balance, charge balance and verification of reaction formulae from the BIGG Models^20^.

### Formulation of biomass composition

The biomass reaction generated in the reconstructed model comprised lipids, proteins, DNA, RNA, cofactors, energy coefficients and cell wall components. The biomass components were determined experimentally to further validate the model as detailed in Supplementary material S1. *A. burnsii* was grown in the log phase. Total RNA and DNA extractions were performed using Spin column fungal total RNA and DNA Purification Kit in triplicates. The RNA and DNA samples were quantified by absorbance using a NanoDrop (Thermo Fisher Scientific, USA). Total protein estimation of fungal cells was performed using the Biuret method, with Bovine Serum Albumin as the standard^21^. The amino acid composition was determined from the *A. burnsii* protein FASTA using the Protein Information Resource (PIR) database. The prevalence of each amino acid was then converted into stoichiometric coefficients (mmol/g DW). The classes of lipids present in *A. burnsii* cells was adapted from the closely related microorganism *Alternaria* sp. MG1^12^. The cell wall content was estimated using the aniline blue fluorescence assay^22^ and quantified using a fluorescence plate reader. Contents of potassium, sodium, and ferrous ions were estimated using the inductively coupled plasma optical emission spectrometry (ICP-OES, Perkin Elmer Optima 5300 DV). The biomass concentration, camptothecin yield, and residual glucose concentration profile in the medium were adapted from the experimental growth kinetics studies^10^. Growth-associated maintenance (GAM) and non-growth-associated maintenance (NGAM) reactions were adapted from *Alternaria* sp. MG1^12^.

### Flux Balance and Flux Scanning Analyses

Flux Balance Analysis (FBA) is a computational method used to determine the flow of metabolites through a metabolic network^23,24^. FBA allows for the prediction of an organism’s growth and production rate of key metabolites. In our study, FBA was applied to analyse our model (*Alt*GEM *i*DD1552) by setting constraints on exchange reactions using experimental data for substrate and metabolite uptake rates with the objective function of maximizing cell biomass. FROG analysis (Flux Variability, Reaction Deletion, Objective Function, and Gene Deletion) was also performed to ensure model reproducibility and quality. Flux Scanning based on Enforced Objective Flux (FSEOF)^25,26^ was utilized to identify and rank targets for over-expression and knockouts to enhance camptothecin production. FSEOF identifies reaction targets with fluxes that increase monotonically when product formation is incrementally enforced while growth is maximized as the objective function. Potential enzyme targets for over-expression and knockouts were ranked based on a score, ‘*f*’, which is a phenotypic fraction weighing reaction fluxes under over-expressed and wildtype conditions^27–29^.

### Culture conditions

*A. burnsii* NCIM 1409 cultures were streaked on slants consisting of Potato Dextrose Agar medium (Himedia, Mumbai) with an initial pH of 5.6 and incubated for 7 days at 28 °C to promote mycelial growth^10^. To prepare the inoculum for initiating suspension cultures for each generation, the spores and mycelia were washed with 5 ml of saline solution (0.9% w/v NaCl). Suspension cultures were initiated by the addition of 2% (v/v) of this inoculum to Erlenmeyer flasks (in triplicates), each containing 50 ml of freshly prepared Potato Dextrose broth medium (Himedia, Mumbai). These cultures were grown for 8 days at 28 °C and 120 rpm. At the end of the growth period, the biomass was harvested and oven-dried for subsequent experiments.

### Cell growth and harvesting for ^13^C isotope labeling experiments

*A. burnsii* slants were initiated in Potato Dextrose Agar (PDA) media at 28 °C. After 7 days of growth, the cells were harvested, washed with saline water and inoculated in the potato infusion broth (Sigma Aldrich, USA) with 2% w/v labelled carbon source. Two different combinations of the labeled carbon source were used for the experiments - 20%[U-^13^C_6_] glucose and 99% [1-^13^C] glucose with four replicates. The cells were harvested on the 8^th^ day, lyophilised and stored at -20 × for further analysis.

### Cell hydrolysis and protein hydrolysate derivatization

The lyophilized cells were acid hydrolysed in 6M HCl and incubated at 105 °C for 24 hours^30^. The hydrolysate volume was adjusted to approximately 500 µl in each tube and centrifuged at 13,000 rpm for 15 minutes. Subsequently, 50 µl of the supernatant was dried using a speed vacuum. Each dried sample was then reconstituted with 60 µl of pyridine and incubated at 37 °C for 30 min at 900 rpm. Next, 100 µl of the derivatizing reagent, MtBSTFA [N-Methyl-N-(t-butyldimethylsilyl trifluoroacetamide)] with 1% tBDMCS (t-butyl dimethylchlorosilane), was added and incubated at 60 °C for 30 min at 900 rpm. The samples were centrifuged at 13,000 rpm for 10 min, and the supernatant was subjected to GC-MS analysis ^31^.

### GC-MS data acquisition and natural ^13^C isotope correction

GC-MS data acquisition was performed using a GC system equipped with an HP-5MS column at the School of Biosciences and Bioengineering, IIT Mandi. Helium served as the carrier gas with a flow rate maintained at 0.6 mL/min. The initial oven temperature was set to 50 °C and held for 5 min. Subsequently, the temperature was increased to 200 °C at a rate of 10 °C/min, with a hold time of 10 min, then further increased to 270 °C at 5 °C/min with a 7-min hold time. Finally, the temperature was raised to 300 °C at 5 °C/min and held for 3 min. An injection volume of 1 µL was used, and the total run time was 60 min. The obtained spectra were baseline-corrected using Metalign software^32^. Amino acid identification was performed with the NIST 17 library and in-house standards to extract retention times and mass ion fragments (m/z). The intensity of each amino acid fragment’s mass ion was determined using Agilent Chemstation software. IsoCor software was utilized to correct the naturally stable isotope abundance from the mass isotopomer distribution (MID) of each fragment ion ^33^. The corrected MID values were further used to calculate the average ^13^C abundance in each fragment, and the valid ones were selected for further analysis^31,34^

## Supporting information

AltGEM_iDD1552

## RESOURCE AVAILABILITY

### Lead contact

Further information and requests for resources and reagents should be directed to and will be fulfilled by the lead contact, Dr. Smita Srivastava

### Materials availability

All materials generated in this study are available from the lead contact with a completed materials transfer agreement.

### Data and code availability

The primary datasets supporting the findings of this study are provided in the supplementary information.

## Acknowledgments

The author(s) declare that financial support was received for the research, authorship, and/or publication of this article. Prof. Smita Srivastava would like to acknowledge the financial support by Higher Education Financing Agency – HEFA (CR2250381BTHEFA008458) and Herbalife International India Private Limited (CR24251461BTHBII008458) for the ongoing research in the Plant Cell Technology Lab at Indian Institute of Technology Madras including the work stated above. Shagun S thanks CSIR-UGC (09/1058(0024)/2020-EMR-I) for the Ph.D. fellowship.

## Author Contribution

Conceptualization, D.D.U.S., S.M. and S.S.; Methodology, D.D.U.S., S.M., and Sh.Sh.; Investigation, D.D.U.S. and S.M; Formal Analysis, D.D.U.S., S.M., and Sh.Sh.; Data Curation, D.D.U.S.; Visualization, S.M.; Writing – Original Draft, D.D.U.S. and S.M.; Writing – Review & Editing, all authors; Supervision, K.R, S.K.M, and S.S.; Funding Acquisition, S.S.

## Declaration of interests

The authors declare no competing interests.

### Supplemental information

Document S1 includes Figures S1–S5, Tables S1-S13, and supplemental references supporting the genome-scale metabolic model reconstruction, biomass composition, and 13C metabolic flux analysis in *Alternaria burnsii* NCIM1409.

## References

1. Kusari, S., Zühlke, S., and Spiteller, M. (2009). An endophytic fungus from Camptotheca acuminata that produces camptothecin and analogues. J Nat Prod 72, 2–7. 10.1021/NP800455B/SUPPL_FILE/NP800455B_SI_001.PDF.

2. Lin, R.K., Ho, C.W., Liu, L.F., and Lyu, Y.L. (2013). Topoisomerase IIβ deficiency enhances camptothecin-induced apoptosis. Journal of Biological Chemistry 288, 7182–7192. 10.1074/JBC.M112.415471/ASSET/6F3388F7-38C8-419C-89AC-E6E6BCA290B9/MAIN.ASSETS/GR5.JPG.

3. Liu, L.F., Desai, S.D., Li, T.K., Mao, Y., Sun, M., and Sim, S.P. (2000). Mechanism of action of camptothecin. Ann. N. Y. Acad. Sci. 922, 1–10. 10.1111/j.1749-6632.2000.tb07020.x.

4. Mathijssen, R.H.J., Verweij, J., Jonge, M.J.A. de, Nooter, K., Stoter, G., and Sparreboom, A. (2002). Impact of Body-Size Measures on Irinotecan Clearance: Alternative Dosing Recommendations. Journal of Clinical Oncology 20, 81–87. 10.1200/JCO.2002.20.1.81.

5. Schulz, B., Boyle, C., Draeger, S., Römmert, A.K., and Krohn, K. (2002). Endophytic fungi: A source of novel biologically active secondary metabolites. Mycol Res 106, 996–1004. 10.1017/S0953756202006342.

6. Gouda, S., Das, G., Sen, S.K., Shin, H.S., and Patra, J.K. (2016). Endophytes: A treasure house of bioactive compounds of medicinal importance. Front Microbiol 7, 219261. 10.3389/FMICB.2016.01538/BIBTEX.

7. Kumari, P., Deepa, N., Trivedi, P.K., Singh, B.K., Srivastava, V., and Singh, A. (2023). Plants and endophytes interaction: a “secret wedlock” for sustainable biosynthesis of pharmaceutically important secondary metabolites. Microbial Cell Factories 2023 22:1 22, 1–19. 10.1186/S12934-023-02234-8.

8. Usman, M., Shah, I.H., Sabir, I.A., Malik, M.S., Rehman, A., Murtaza, G., Azam, M., Rahman S. ur, Rehman, A., Ashraf, G.A., et al. (2024). Synergistic partnerships of endophytic fungi for bioactive compound production and biotic stress management in medicinal plants. Plant Stress 11, 100425. 10.1016/J.STRESS.2024.100425.

9. Hashem, A.H., Attia, M.S., Kandil, E.K., Fawzi, M.M., Abdelrahman, A.S., Khader, M.S., Khodaira, M.A., Emam, A.E., Goma, M.A., and Abdelaziz, A.M. (2023). Bioactive compounds and biomedical applications of endophytic fungi: a recent review. Microbial Cell Factories 2023 22:1 22, 1–23. 10.1186/S12934-023-02118-X.

10. Mohinudeen, I.A.H.K., Kanumuri, R., Soujanya, K.N., Shaanker, R.U., Rayala, S.K., and Srivastava, S. (2021). Sustainable production of camptothecin from an Alternaria sp. isolated from Nothapodytes nimmoniana. Scientific Reports 2021 11:1 11, 1–11. 10.1038/s41598-020-79239-5.

11. DiCenzo, G.C., Checcucci, A., Bazzicalupo, M., Mengoni, A., Viti, C., Dziewit, L., Finan, T.M., Galardini, M., and Fondi, M. (2016). Metabolic modelling reveals the specialization of secondary replicons for niche adaptation in Sinorhizobium meliloti. Nature Communications 2016 7:1 7, 1–10. 10.1038/ncomms12219.

12. Lu, Y., Ye, C., Che, J., Xu, X., Shao, D., Jiang, C., Liu, Y., and Shi, J. (2019). Genomic sequencing, genome-scale metabolic network reconstruction, and in silico flux analysis of the grape endophytic fungus Alternaria sp. MG1. Microb Cell Fact 18, 1–16. 10.1186/S12934-019-1063-7/TABLES/4.

13. Kanehisa, M., Furumichi, M., Sato, Y., Matsuura, Y., and Ishiguro-Watanabe, M. (2025). KEGG: biological systems database as a model of the real world. Nucleic Acids Res 53, D672–D677. 10.1093/NAR/GKAE909.

14. BioCyc Pathway/Genome Database Collection https://www.biocyc.org/.

15. MetaCyc: Metabolic Pathways From all Domains of Life https://metacyc.org/.

16. Saier, M.H., Tran, C. V., and Barabote, R.D. (2006). TCDB: the Transporter Classification Database for membrane transport protein analyses and information. Nucleic Acids Res 34, D181–D186. 10.1093/NAR/GKJ001.

17. ModelSEED https://modelseed.org/genomes/.

18. Heirendt, L., Arreckx, S., Pfau, T., Mendoza, S.N., Richelle, A., Heinken, A., Haraldsdóttir, H.S., Wachowiak, J., Keating, S.M., Vlasov, V., et al. (2019). Creation and analysis of biochemical constraint-based models using the COBRA Toolbox v.3.0. Nature Protocols 2019 14:3 14, 639– 702. 10.1038/s41596-018-0098-2.

19. Hawkins, C., Ginzburg, D., Zhao, K., Dwyer, W., Xue, B., Xu, A., Rice, S., Cole, B., Paley, S., Karp, P., et al. (2021). Plant Metabolic Network: A multi-species resource of plant metabolic information. bioRxiv, 2021.03.30.437738. 10.1101/2021.03.30.437738.

20. King, Z.A., Lu, J., Dräger, A., Miller, P., Federowicz, S., Lerman, J.A., Ebrahim, A., Palsson, B.O., and Lewis, N.E. (2016). BiGG Models: A platform for integrating, standardizing and sharing genome-scale models. Nucleic Acids Res 44, D515–D522. 10.1093/NAR/GKV1049.

21. Parvin, R., Pande, S. V., and Venkitasubramanian, T.A. (1965). On the colorimetric biuret method of protein determination. Anal Biochem 12, 219–229. 10.1016/0003-2697(65)90085-0.

22. Smith, M.M., and McCully, M.E. (1978). Enhancing Aniline Blue Fluorescent Staining of Cell Wall Structures. Stain Technol 53, 79–85. 10.3109/10520297809111446.

23. Orth, J.D., Thiele, I., and Palsson, B.O. (2010). What is flux balance analysis? Nat Biotechnol 28, 245–248. 10.1038/NBT.1614.

24. Raman, K., and Chandra, N. (2009). Flux balance analysis of biological systems: applications and challenges. Brief Bioinform 10, 435–449. 10.1093/BIB/BBP011.

25. Alper, H., Jin, Y.S., Moxley, J.F., and Stephanopoulos, G. (2005). Identifying gene targets for the metabolic engineering of lycopene biosynthesis in Escherichia coli. Metab Eng 7, 155–164. 10.1016/J.YMBEN.2004.12.003.

26. Choi, H.S., Lee, S.Y., Kim, T.Y., and Woo, H.M. (2010). In silico identification of gene amplification targets for improvement of lycopene production. Appl Environ Microbiol 76, 3097–3105. 10.1128/AEM.00115-10/SUPPL_FILE/SUPPLEMENTAL7SUPPLEMENTAL_FIGURE_LEGENDS.DOC.

27. Raajaraam, L., and Raman, K. (2022). A Computational Framework to Identify Metabolic Engineering Strategies for the Co-Production of Metabolites. Front Bioeng Biotechnol 9. 10.3389/FBIOE.2021.779405.

28. Boghigian, B.A., Armando, J., Salas, D., and Pfeifer, B.A. (2012). Computational identification of gene over-expression targets for metabolic engineering of taxadiene production. Appl Microbiol Biotechnol 93, 2063–2073. 10.1007/S00253-011-3725-1.

29. Murali, S., Ibrahim, M., Rajendran, H., Shagun, S., Masakapalli, S.K., Raman, K., and Srivastava, S. (2023). Genome-scale metabolic model led engineering of Nothapodytes nimmoniana plant cells for high camptothecin production. Front Plant Sci 14, 1207218. 10.3389/FPLS.2023.1207218/BIBTEX.

30. Schwechheimer, S.K., Becker, J., Peyriga, L., Portais, J.C., Sauer, D., Müller, R., Hoff, B., Haefner, S., Schröder, H., Zelder, O., et al. (2018). Improved riboflavin production with Ashbya gossypii from vegetable oil based on 13C metabolic network analysis with combined labeling analysis by GC/MS, LC/MS, 1D, and 2D NMR. Metab Eng 47, 357–373. 10.1016/J.YMBEN.2018.04.005.

31. Jyoti, P., Shree, M., Joshi, C., Prakash, T., Ray, S.K., Satapathy, S.S., and Masakapalli, S.K. (2020). The Entner-Doudoroff and Nonoxidative Pentose Phosphate Pathways Bypass Glycolysis and the Oxidative Pentose Phosphate Pathway in Ralstonia solanacearum. mSystems 5. 10.1128/MSYSTEMS.00091-20/SUPPL_FILE/MSYSTEMS.00091-20-SF003.TIF.

32. Lommen, A., and Kools, H.J. (2012). MetAlign 3.0: Performance enhancement by efficient use of advances in computer hardware. Metabolomics 8, 719–726. 10.1007/S11306-011-0369-1/FIGURES/3.

33. Millard, P., Letisse, F., Sokol, S., and Portais, J.C. (2012). IsoCor: correcting MS data in isotope labeling experiments. Bioinformatics 28, 1294–1296. 10.1093/BIOINFORMATICS/BTS127.

34. Shree, M., and Masakapalli, S.K. (2018). Intracellular Fate of Universally Labelled 13C Isotopic Tracers of Glucose and Xylose in Central Metabolic Pathways of Xanthomonas oryzae. Metabolites 2018, Vol. 8, Page 66 8, 66. 10.3390/METABO8040066.

35. Natarajan, S., Pucker, B., and Srivastava, S. (2023). Genomic and transcriptomic analysis of camptothecin producing novel fungal endophyte: Alternaria burnsii NCIM 1409. Scientific Reports 2023 13:1 13, 1–11. 10.1038/s41598-023-41738-6.

36. Wattam, A.R., Abraham, D., Dalay, O., Disz, T.L., Driscoll, T., Gabbard, J.L., Gillespie, J.J., Gough, R., Hix, D., Kenyon, R., et al. (2014). PATRIC, the bacterial bioinformatics database and analysis resource. Nucleic Acids Res 42, D581–D591. 10.1093/NAR/GKT1099.

37. Buldum, G., Bismarck, A., and Mantalaris, A. (2018). Recombinant biosynthesis of bacterial cellulose in genetically modified Escherichia coli. Bioprocess Biosyst Eng 41, 265–279. 10.1007/s00449-017-1864-1.

38. Rather, G.A., Sharma, A., Misra, P., Kumar, A., Kaul, V., and Lattoo, S.K. (2020). Molecular characterization and overexpression analyses of secologanin synthase to understand the regulation of camptothecin biosynthesis in Nothapodytes nimmoniana (Graham.) Mabb. Protoplasma 257, 391–405. 10.1007/S00709-019-01440-9/FIGURES/7.

39. Sharma, A., Verma, P., Mathur, A., and Mathur, A.K. (2018). Overexpression of tryptophan decarboxylase and strictosidine synthase enhanced terpenoid indole alkaloid pathway activity and antineoplastic vinblastine biosynthesis in Catharanthus roseus. Protoplasma 255, 1281–1294. 10.1007/S00709-018-1233-1.

40. F.M., L., Currin, A., and Dixon, N. (2019). Directed evolution of the PcaV allosteric transcription factor to generate a biosensor for aromatic aldehydes. J. Biol. Eng. 13. 10.1186/s13036-019-0214-z.

41. Chen, Y., Wang, Y., Liu, M., Qu, J., Yao, M., Li, B., Ding, M., Liu, H., Xiao, W., and Yuan, Y. (2019). Primary and secondary metabolic effects of a key gene deletion (ΔYPL062W) in metabolically engineered terpenoid-producing saccharomyces cerevisiae. Appl Environ Microbiol 85. 10.1128/AEM.01990-18/SUPPL_FILE/AEM.01990-18-S0001.PDF.

42. Özaydin, B., Burd, H., Lee, T.S., and Keasling, J.D. (2013). Carotenoid-based phenotypic screen of the yeast deletion collection reveals new genes with roles in isoprenoid production. Metab Eng 15, 174–183. 10.1016/J.YMBEN.2012.07.010.

43. Schadeweg, V., and Boles, E. (2016). N-Butanol production in Saccharomyces cerevisiae is limited by the availability of coenzyme A and cytosolic acetyl-CoA. Biotechnol Biofuels 9, 1–12. 10.1186/S13068-016-0456-7/FIGURES/5.

44. Lian, J., Si, T., Nair, N.U., and Zhao, H. (2014). Design and construction of acetyl-CoA overproducing Saccharomyces cerevisiae strains. Metab Eng 24, 139–149. 10.1016/J.YMBEN.2014.05.010.

45. Jouhten, P., Pitkänen, E., Pakula, T., Saloheimo, M., Penttilä, M., and Maaheimo, H. (2009). 13C-metabolic flux ratio and novel carbon path analyses confirmed that Trichoderma reesei uses primarily the respirative pathway also on the preferred carbon source glucose. BMC Syst Biol 3, 104. 10.1186/1752-0509-3-104/FIGURES/6.

46. Schwechheimer, S.K., Becker, J., Peyriga, L., Portais, J.C., Sauer, D., Müller, R., Hoff, B., Haefner, S., Schröder, H., Zelder, O., et al. (2018). Improved riboflavin production with Ashbya gossypii from vegetable oil based on 13C metabolic network analysis with combined labeling analysis by GC/MS, LC/MS, 1D, and 2D NMR. Metab Eng 47, 357–373. 10.1016/J.YMBEN.2018.04.005.

