## Supplementary File for "Genome-scale metabolic model guided metabolic flux analysis in the endophyte *Alternaria burnsii* NCIM1409"

[1] Department of Biotechnology, Bhupat and Jyoti Mehta School of Biosciences, Indian Institute of Technology Madras
[2] The Novo Nordisk Foundation Center for Biosustainability, Technical University of Denmark, Kongens Lyngby, 2800, Denmark
[3] Herbalife-IITM Plant Cell Fermentation Technology Lab, Indian Institute of Technology Madras Research Park, Chennai, 600113, India
[4] School of Biosciences and Bioengineering, Indian Institute of Technology Mandi, Kamand 175075, India
[5] Department of Data Science and AI, Wadhwani School of Data Science and AI, Indian Institute of Technology Madras, Chennai, 600 036, India
[6] Robert Bosch Centre for Data Science and Artificial Intelligence (RBCDSAI), Indian Institute of Technology Madras, Chennai, 600 036, India
[7] Lead contact
*equally contributed

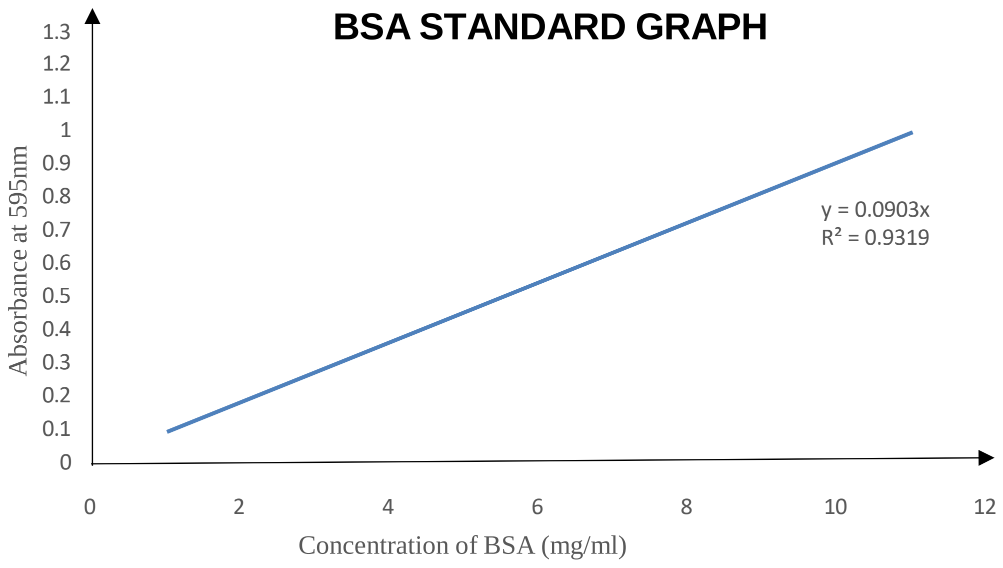


***Figure S1:*** *The standard curve of BSA obtained by plotting various concentrations of BSA for estimation of protein concentration in Alternaria burnsii.*

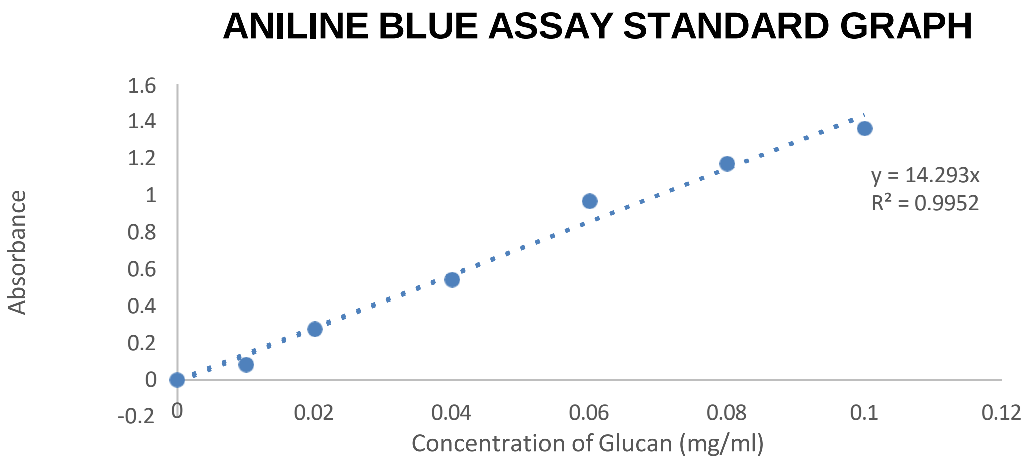


***Figure S2*** *The standard curve of aniline blue assay obtained by plotting various concentrations of glucan for estimation of cell wall concentration in Alternaria burnsii*

The composition of DNA was based on the genomic sequence of *Alternaria burnsii*

G+C content (%): 43.57

| DNA | | | | | |
|---|---|---|---|---|---|
| G+C content (%): 43.57 | | | | | |
| **DNA** | **Prevalence (%)** | **MW (g/mol)** | **P*MW (%)** | **by weight (%)** | **mmol/gDW** |
| dATP | 50 | 487.00 | 13746.81 | 28.47 | 0.00585 |
| dCTP | 0 | 462.99 | 10084.34 | 20.88 | 0.00451 |
| dGTP | 0 | 503.00 | 10960.56 | 22.71 | 0.00451 |
| dTTP | 50 | 477.99 | 13486.49 | 27.94 | 0.00585 |
| **Total** | **100** | | **48278.21** | 100.00 | |

***Table S1*** *DNA composition*

| RNA | | | | | |
|---|---|---|---|---|---|
| **RNA** | **Prevalence (%)** | **MW (g/mol)** | **P*MW (%)** | **by weight (%)** | **mmol/gDW** |
| ATP | 25.60 | 503.00 | 128.77 | 25.92 | 0.01185 |
| CTP | 19.60 | 478.98 | 93.88 | 18.90 | 0.00907 |
| GTP | 28.60 | 518.99 | 148.43 | 29.88 | 0.01324 |
| UTP | 26.20 | 479.97 | 125.75 | 25.31 | 0.01213 |
| **Total** | **100** | | **496.83** | 100.00 | |

***Table S2*** *RNA composition*

| Lipids (Adapted from *Alternaria* sp. MG1) | | | | | |
|---|---|---|---|---|---|
| **Components** | **Content % (w/w)** | **MW (g/mol)** | **P*MW (%)** | **by weight (%)** | **mmol/gDCW** |
| Triacylglycerol | 31.0 | 955.0 | 29603.8 | 6.64 | 0.01555 |
| Free fatty acids | 5.0 | 301.3 | 1506.6 | 0.34 | 0.00251 |
| Phosphatidylethanolamine | 14.0 | 782.5 | 10955.0 | 2.46 | 0.00702 |
| Phosphatidylcholine | 35.0 | 834.8 | 29218.0 | 6.56 | 0.01755 |
| Phosphatidylserine | 6.0 | 827.3 | 4963.9 | 1.11 | 0.00301 |
| Phosphatidylamine | 9.0 | 755.2 | 6797.2 | 1.53 | 0.00451 |
| **Total** | **100.0** | **4456.1** | 445613.0 | | |

***Table S3*** *Phospholipids composition*

| Glycogen | | | | | |
| --- | --- | --- | --- | --- | --- |
| **Components** | **Content % (w/w) *** | **MW (g/mol)** | **P*MW (%)** | **by weight (%)** | **mmol/gDCW** |
| Glycogen | 100 | 666.6 | 666.6 | 36837.59398 | 0.00150 |
| | **100.0** | **666.6** | **666.6** | **36837.6** | |

***Table S4*** *Glycogen composition*

| Cell Wall | | | | | |
| --- | --- | --- | --- | --- | --- |
| **Components** | **Content % (w/w) *** | **MW (g/mol)** | **P*MW (%)** | **% (mole)** | **mmol/gDCW** |
| chitin | 27 | 203.2 | 54.864 | 100 | 0.26575 |
| glucan | 73 | 162.1 | 118.333 | | 0.90068 |
| | **100.0** | | **173.2** | | |

***Table S5*** *Carbohydrates composition (Adapted from Alternaria sp. MG1)*

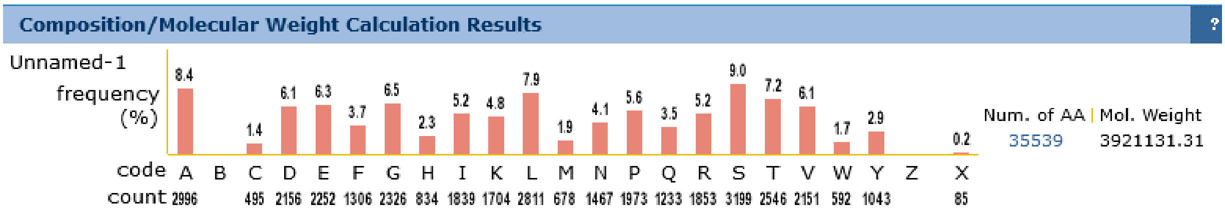


***Figure S3*** *The amino acids were identified in this study from PIR database*

| Amino Acids | Abbreviation | Letter Code | Prevalence (% | MW (g/mol) | P*MW (%) | By weight (%) | mmol/gDW |
|---|---|---|---|---|---|---|---|
| **Alanine (A)** | Ala | A | 8.4 | 89.05 | 748.02 | 6.206 | 0.0985 |
| **Arginine (R)** | Arg | R | 5.2 | 175.11 | 910.572 | 7.5546 | 0.0423 |
| **Asparagine (N)** | Asn | N | 4.1 | 132.05 | 541.405 | 4.4918 | 0.1842 |
| **Aspartic acid (D)** | Asp | D | 6.1 | 132.04 | 805.444 | 6.6824 | 0.094 |
| **Cysteine (C)** | Cys | C | 1.4 | 121.02 | 169.428 | 1.4057 | 0.0599 |
| **Glutamate (E)** | Gln | Q | 3.5 | 146.05 | 511.175 | 4.241 | 0.1118 |
| **Glutamine (Q)** | Glu | E | 6.3 | 146.07 | 920.241 | 7.6348 | 0.1527 |
| **Glycine (G)** | Gly | G | 6.5 | 75.03 | 487.695 | 4.0462 | 0.0846 |
| **Histidine (H)** | His | H | 2.3 | 155.07 | 356.661 | 2.9591 | 0.0293 |
| **Isoleucine (I)** | Ile | I | 5.2 | 131.09 | 681.668 | 5.6555 | 0.0454 |
| **Leucine (L)** | Leu | L | 7.9 | 131.09 | 1035.611 | 8.592 | 0.0738 |
| **Lysine (K)** | Lys | K | 4.8 | 147.11 | 706.128 | 5.8584 | 0.0683 |
| **Methionine (M)** | Met | M | 1.9 | 149.05 | 283.195 | 2.3495 | 0.119 |
| **Phenylalanine (F)** | Phe | F | 3.7 | 165.08 | 610.796 | 5.0675 | 0.0372 |
| **Proline (P)** | Pro | P | 5.6 | 115.06 | 644.336 | 5.3458 | 0.0385 |
| **Serine (S)** | Ser | S | 9 | 105.04 | 945.36 | 7.8432 | 0.0672 |
| **Threonine (T)** | Thr | T | 7.2 | 119.06 | 857.232 | 7.1121 | 0.0627 |
| **Tryptophan (W)** | Trp | W | 1.7 | 204.09 | 346.953 | 2.8785 | 0.0705 |
| **Tyrosine (Y)** | Tyr | Y | 2.9 | 181.07 | 525.103 | 4.3565 | 0.0334 |
| **Valine (V)** | Val | V | 6.1 | 117.08 | 714.188 | 5.9253 | 0.0465 |
| | | **Total** | 100 | | **12053.19** | | |

***Table S6*** *Protein composition*

| Glycerol | | | | | |
|---|---|---|---|---|---|
| **Components** | **Content % (w/w)** | **MW (g/mol)** | **P*MW (%)** | **by weight (%)** | **mmol/gDCW** |
| glycerol | 100 | 92.1 | 92.1 | 100 | 0.06515 |

***Table S7*** *Glycerol (Adapted from Alternaria sp. MG1)*

| D-Mannitol | | | | | |
|---|---|---|---|---|---|
| **Components** | **Content % (w/w)** | **MW (g/mol)** | **P*MW (%)** | **by weight (%)** | **mmol/gDCW** |
| D-mannitol | 100 | 182.2 | 182.2 | 100 | 0.33480 |

***Table S8*** *Mannitol (Adapted from Alternaria sp. MG1)*

| ICP OES Analysis | | | |
|---|---|---|---|
| **Element** | **Average Concentration (mg/L)** | **Molecular weight (g/mol)** | **mmol/g** |
| Fe 238.204 | 0.0413 | 55.845 | 450.7264 |
| K 766.490 | 2.693 | 39.0983 | 4.839497 |
| Na 589.592 | 17.753 | 22.9897 | 4.316585 |

***Table S9*** *Elemental Ion concentration*

**Biomass equation:** 71.4 h2o[c] + 71.4 atp[c] + 0.0985 ala[c] + 0.0423 arg[c] + 0.1842 asn[c] + 0.094 asp[c] + 0.0599 cys[c] + 0.1118 glu[c] + 0.1527 gln[c] + 0.0846 gly[c] +0.0293 his[c] + 0.0454 ile[c] + 0.0738 leu[c] + 0.0683 lys[c] + 0.119 met[c] +0.0372 phe[c] + 0.0385 pro[c] + 0.0672 ser[c] + 0.0627 thr[c] + 0.0705 trp[c] + 0.0334 tyr[c] + 0.0465 val[c] + 0.00585 damp[c] + 0.00451 dgmp[c] + 0.00451 dcmp[c] + 0.00585 dtmp[c] + 0.0234 amp[c] + 0.0237 gmp[c] + 0.0222 cmp[c] + 0.0225 ump[c] + 0.020289 egstr[c] + 0.45310 chit[c] + 1.53566 13glucan[c] + 0.11526 mnt[c] + 0.002333 pins[c] + 0.001676 cl[c] + 0.0155 tagly[c] + 0.00702 pe[c] + 0.00301 ps[c] + 0.00301 pg[c] + 0.01755 pc[c] + 0.00451 pa[c] -> 71.4 h[c] + 71.4 pi[c] + 71.4 adp[c]

| CELLULAR COMPOSITION | | |
|---|---|---|
| **Items** | **Composition (% w/w)** | **Reference** |
| Protein | 36.80 | This study |
| DNA | 0.80 | |
| RNA | 2.30 | |
| Lipid | 22.35 | |
| Glycogen | 0.10 | (1) |
| Cell wall components (Chitin and Glucan) | 34.10 | |
| Ash content | 1.00 | |
| Glycerol | 0.60 | |
| D-mannitol | 2.10 | |
| **Total** | **100** | |

***Table S10*** *Biomass composition of Alternaria burnsii*

| **Substrate** | ***Constraint*** | ***Specific Growth rate ($h^{-1}$)*** | |
|---|---|---|---|
| | Maximum Substrate Uptake/ Utilization rate (mmol $gDW^{-1}$ $h^{-1}$) | Model | Experimental |
| Glucose | 1.9274 | 0.022 | 0.028 |
| Fructose | 1.9274 | 0.015 | 0.016 |
| Sucrose | 1.0144 | 0.009 | 0.009 |
| Lactose | 1.0144 | 0.005 | 0.004 |
| Maltose | 1.0144 | 0.01 | 0.009 |
| Galactose | 1.9274 | 0.018 | 0.019 |
| Cellulose | 2.1415 | 0.012 | 0.011 |
| Xylose | 2.3128 | 0.01 | 0.009 |
| Starch | 0.2964 | 0.0043 | 0.0045 |
| Mannitol | 1.906 | 0.0023 | 0.0026 |

***Table S11*** *Comparison between model and experimental values for specific growth rate.*

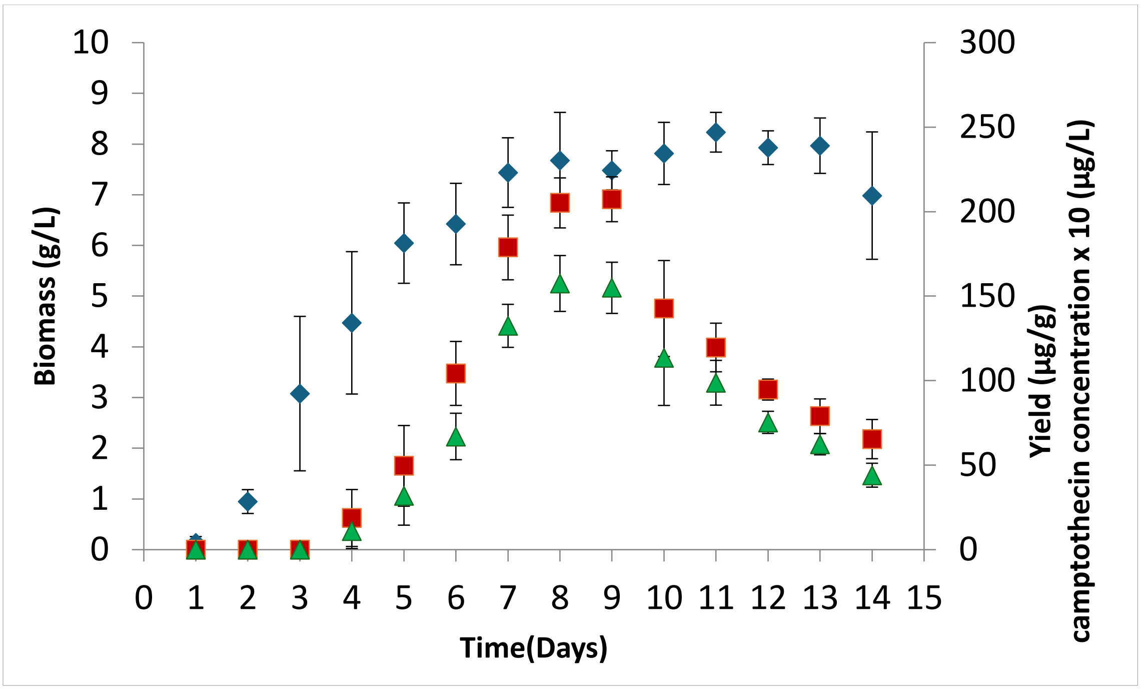


***Figure S4*** *Time-course analysis of biomass accumulation and camptothecin production. Biomass (g/L) is represented by blue diamonds (◊), measured on the left y-axis. Camptothecin yield (µg/g dry cell weight) is shown by red squares (■), and camptothecin concentration in culture (µg/L, scaled ×10) is indicated by green triangles (▲), both plotted on the right y-axis. Data represent the mean ± standard deviation of three biological replicates. Maximum biomass was observed around day 9, while camptothecin production peaked earlier and gradually declined after day 10.(2)*

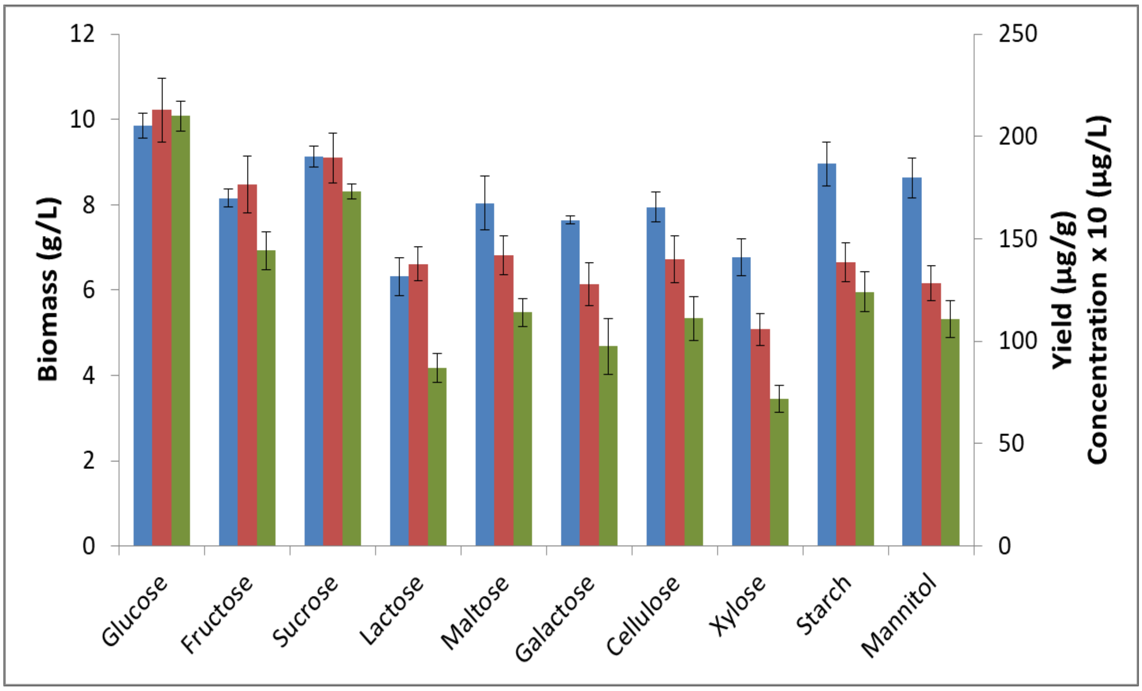


***Figure S5*** *Effect of different carbon sources on biomass production and camptothecin yield. Biomass concentration (g/L) is shown by blue bars and plotted on the left y-axis. Camptothecin yield (µg/g dry cell weight) and camptothecin concentration in culture (µg/L, scaled ×10) are represented by red and green bars, respectively, and plotted on the right y-axis. Glucose supported the highest biomass and camptothecin production, while alternative carbon sources such as xylose, starch, and mannitol showed*

*comparatively lower values. Data are presented as mean ± standard deviation (n = 3). Previous Study taken from (2)*

| Amino acid fragment | Fragment ion | Carbon number | 99% $^{12}C$ glucose | 99% [1-$^{13}C$] glucose | 20%[U-$^{13}C_6$] glucose |
|---|---|---|---|---|---|
| Ala260 | $[M-57]^+$ | C1-C3 | 0.91 ± 0.36 | 4.94 ± 0.50 | 3.07 ± 0.30 |
| Ala232 | $[M-85]^+$ | C2-C3 | 1.32 ± 0.38 | 6.04 ± 0.67 | 3.51 ± 0.22 |
| Asp418 | $[M-57]^+$ | C1-C4 | 0.05 ± 0.10 | 5.36 ± 1.32 | 1.74 ± 0.26 |
| Asp390 | $[M-85]^+$ | C2-C4 | 0.06 ± 0.08 | 5.90 ± 1.26 | 1.56 ± 0.20 |
| Glu432 | $[M-57]^+$ | C1-C5 | 0.17 ± 0.19 | 5.53 ± 0.44 | 2.22 ± 0.46 |
| Glu330 | $[M-85]^+$ | C2-C5 | 0.03 ± 0.03 | 5.92 ± 0.56 | 2.08 ± 0.18 |
| Gly246 | $[M-57]^+$ | C1-C3 | 1.22 ± 0.41 | 2.00 ± 0.27 | 2.01 ± 0.38 |
| Gly218 | $[M-85]^+$ | C2-C3 | 2.15 ± 0.54 | 2.53 ± 1.01 | 2.75 ± 0.37 |
| His440 | [M-57]+ | C1-C6 | 0.04 ± 0.03 | 1.37 ± 0.41 | 0.70 ± 0.18 |
| His412 | $[M-85]^+$ | C2-C6 | 0.00 ± 0.00 | 0.71 ± 0.50 | 0.28 ± 0.34 |
| Ile302 | $[M-57]^+$ | C1-C6 | 0.23 ± 0.14 | 2.47 ± 0.27 | 1.36 ± 0.30 |
| Ile274 | $[M-85]^+$ | C2-C6 | 0.29 ± 0.19 | 3.70 ± 0.36 | 3.90 ± 3.99 |
| Leu302 | $[M-57]^+$ | C1-C6 | 0.39 ± 0.13 | 1.79 ± 0.42 | 0.81 ± 0.18 |
| Leu274 | [M-85]+ | C2-C6 | 0.56 ± 0.08 | 2.18 ± 0.62 | 0.93 ± 0.18 |
| Lys431 | $[M-57]^+$ | C1-C6 | 0.22 ± 0.27 | 1.70 ± 0.65 | 0.25 ± 0.29 |
| Phe336 | $[M-57]^+$ | C1-C9 | 0.21 ± 0.05 | 1.78 ± 0.80 | 0.64 ± 0.30 |
| Phe308 | $[M-85]^+$ | C2-C9 | 0.20 ± 0.06 | 1.87 ± 0.73 | 0.78 ± 0.21 |
| Pro286 | $[M-57]^+$ | C1-C5 | 0.08 ± 0.02 | 3.42 ± 0.49 | 1.14 ± 0.39 |
| Pro258 | $[M-85]^+$ | C2-C5 | 0.25 ± 0.07 | 3.69 ± 0.63 | 1.50 ± 0.30 |
| Ser390 | $[M-57]^+$ | C1-C3 | 0.39 ± 0.18 | 2.55 ± 0.51 | 1.22 ± 0.20 |
| Ser262 | $[M-85]^+$ | C2-C3 | 0.14 ± 0.21 | 3.16 ± 0.57 | 1.04 ± 0.26 |
| Thr404 | $[M-57]^+$ | C1-C4 | 0.19 ± 0.06 | 4.24 ± 0.31 | 1.91 ± 0.35 |
| Thr376 | $[M-85]^+$ | C2-C4 | 0.07 ± 0.08 | 4.78 ± 0.39 | 1.74 ± 0.39 |
| Tyr466 | $[M-57]^+$ | C1-C9 | 0.00 ± 0.00 | 2.06 ± 0.63 | 0.90 ± 0.35 |
| Val288 | $[M-57]^+$ | C1-C5 | 0.38 ± 0.05 | 2.76 ± 0.57 | 1.57 ± 0.18 |
| Val260 | $[M-85]^+$ | C2-C5 | 0.77 ± 0.16 | 3.76 ± 0.61 | 2.20 ± 0.36 |

***Table S12*** *Average $^{13}C$ abundance [in %age, represented as Mean ± Standard deviation (SD); n=4] in valid amino acid fragments ($[M-57]^+$ and $[M-85]^+$) of Alternaria burnsii grown in Potato infusion broth supplemented with 99% [$^{12}C$]glucose, 99 % [1-$^{13}C$]glucose and 20% [U-$^{13}C_6$]glucose [Ala: Alanine; Asp: Aspartic acid; Glu: Glutamic acid; Gly: Glycine; His: Histidine; Ile: Isoleucine; Leu: Leucine; Lys: Lysine; Phe: Phenylalanine; Pro: Proline; Ser: Serine; Thr: Threonine; Tyr: Tyrosine; Val: Valine].*

| Metabolite fragment | Fragment ion | Carbon number | Mass Isotopomers | 99% $^{12}C$ glucose | 20%[U-$^{13}C_6$] glucose | 99% [1-$^{13}C$] glucose |
|---|---|---|---|---|---|---|
| Ala260 | $[M-57]^+$ | C1-C3 | M+0 | 0.98 ± 0.01 | 0.94 ± 0.01 | 0.86 ± 0.01 |
| | | | M+1 | 0.02 ± 0.01 | 0.03 ± 0.00 | 0.14 ± 0.01 |
| | | | M+2 | 0.00 ± 0.00 | 0.01 ± 0.00 | 0.01 ± 0.00 |
| | | | M+3 | 0.00 ± 0.00 | 0.01 ± 0.00 | 0.00 ± 0.00 |
| Ala232 | $[M-85]^+$ | C2-C3 | M+0 | 0.98 ± 0.01 | 0.95 ± 0.00 | 0.88 ± 0.01 |
| | | | M+1 | 0.02 ± 0.00 | 0.03 ± 0.00 | 0.12 ± 0.01 |
| | | | M+2 | 0.00 ± 0.00 | 0.02 ± 0.00 | 0.00 ± 0.00 |
| Asp418 | $[M-57]^+$ | C1-C4 | M+0 | 1.00 ± 0.00 | 0.95 ± 0.01 | 0.80 ± 0.04 |
| | | | M+1 | 0.00 ± 0.00 | 0.04 ± 0.01 | 0.19 ± 0.03 |
| | | | M+2 | 0.00 ± 0.00 | 0.01 ± 0.01 | 0.01 ± 0.01 |
| | | | M+3 | 0.00 ± 0.00 | 0.00 ± 0.00 | 0.00 ± 0.00 |
| | | | M+4 | 0.00 ± 0.00 | 0.00 ± 0.00 | 0.00 ± 0.00 |
| Asp390 | $[M-85]^+$ | C2-C4 | M+0 | 1.00 ± 0.00 | 0.96 ± 0.01 | 0.83 ± 0.03 |
| | | | M+1 | 0.00 ± 0.00 | 0.03 ± 0.01 | 0.16 ± 0.03 |
| | | | M+2 | 0.00 ± 0.00 | 0.00 ± 0.00 | 0.01 ± 0.00 |
| | | | M+3 | 0.00 ± 0.00 | 0.00 ± 0.00 | 0.00 ± 0.00 |
| Glu432 | $[M-57]^+$ | C1-C5 | M+0 | 0.99 ± 0.01 | 0.92 ± 0.01 | 0.75 ± 0.02 |
| | | | M+1 | 0.01 ± 0.01 | 0.06 ± 0.01 | 0.22 ± 0.02 |
| | | | M+2 | 0.00 ± 0.00 | 0.02 ± 0.01 | 0.03 ± 0.01 |
| | | | M+3 | 0.00 ± 0.00 | 0.00 ± 0.00 | 0.00 ± 0.00 |
| | | | M+4 | 0.00 ± 0.00 | 0.00 ± 0.00 | 0.00 ± 0.00 |
| | | | M+5 | 0.00 ± 0.00 | 0.00 ± 0.00 | 0.00 ± 0.00 |
| Glu330 | $[M-85]^+$ | C2-C5 | M+0 | 1.00 ± 0.00 | 0.94 ± 0.01 | 0.78 ± 0.02 |
| | | | M+1 | 0.00 ± 0.00 | 0.05 ± 0.00 | 0.21 ± 0.01 |
| | | | M+2 | 0.00 ± 0.00 | 0.02 ± 0.00 | 0.02 ± 0.00 |
| | | | M+3 | 0.00 ± 0.00 | 0.00 ± 0.00 | 0.00 ± 0.00 |
| | | | M+4 | 0.00 ± 0.00 | 0.00 ± 0.00 | 0.00 ± 0.00 |
| Gly246 | $[M-57]^+$ | C1-C3 | M+0 | 0.98 ± 0.01 | 0.97 ± 0.01 | 0.96 ± 0.01 |
| | | | M+1 | 0.02 ± 0.01 | 0.03 ± 0.00 | 0.04 ± 0.01 |
| | | | M+2 | 0.00 ± 0.00 | 0.01 ± 0.00 | 0.00 ± 0.00 |
| Gly218 | $[M-85]^+$ | C2-C3 | M+0 | 0.98 ± 0.01 | 0.97 ± 0.00 | 0.97 ± 0.01 |
| | | | M+1 | 0.02 ± 0.01 | 0.03 ± 0.00 | 0.03 ± 0.01 |
| His440 | $[M-57]^+$ | C1-C6 | M+0 | 1.00 ± 0.00 | 0.98 ± 0.01 | 0.92 ± 0.02 |
| | | | M+1 | 0.00 ± 0.00 | 0.01 ± 0.01 | 0.08 ± 0.02 |
| | | | M+2 | 0.00 ± 0.00 | 0.00 ± 0.00 | 0.00 ± 0.00 |
| | | | M+3 | 0.00 ± 0.00 | 0.01 ± 0.00 | 0.00 ± 0.00 |
| | | | M+4 | 0.00 ± 0.00 | 0.00 ± 0.00 | 0.00 ± 0.00 |
| | | | M+5 | 0.00 ± 0.00 | 0.00 ± 0.00 | 0.00 ± 0.00 |
| | | | M+6 | 0.00 ± 0.00 | 0.00 ± 0.00 | 0.00 ± 0.00 |
| His412 | $[M-85]^+$ | C2-C6 | M+0 | 1.00 ± 0.00 | 0.99 ± 0.02 | 0.96 ± 0.02 |
| | | | M+1 | 0.00 ± 0.00 | 0.01 ± 0.02 | 0.04 ± 0.02 |
| | | | M+2 | 0.00 ± 0.00 | 0.00 ± 0.00 | 0.00 ± 0.00 |

| | | | | | | |
|---|---|---|---|---|---|---|
| | | | M+3 | 0.00 ± 0.00 | 0.00 ± 0.00 | 0.00 ± 0.00 |
| | | | M+4 | 0.00 ± 0.00 | 0.00 ± 0.00 | 0.00 ± 0.00 |
| | | | M+5 | 0.00 ± 0.00 | 0.00 ± 0.00 | 0.00 ± 0.00 |
| Ile302 | $[M-57]^+$ | C1-C6 | M+0 | 0.99 ± 0.01 | 0.94 ± 0.01 | 0.87 ± 0.01 |
| | | | M+1 | 0.01 ± 0.01 | 0.04 ± 0.01 | 0.12 ± 0.01 |
| | | | M+2 | 0.00 ± 0.00 | 0.02 ± 0.01 | 0.01 ± 0.00 |
| | | | M+3 | 0.00 ± 0.00 | 0.00 ± 0.00 | 0.00 ± 0.00 |
| | | | M+4 | 0.00 ± 0.00 | 0.00 ± 0.00 | 0.00 ± 0.00 |
| | | | M+5 | 0.00 ± 0.00 | 0.00 ± 0.00 | 0.00 ± 0.00 |
| | | | M+6 | 0.00 ± 0.00 | 0.00 ± 0.00 | 0.00 ± 0.00 |
| Ile274 | $[M-85]^+$ | C2-C6 | M+0 | 0.99 ± 0.01 | 0.88 ± 0.10 | 0.83 ± 0.01 |
| | | | M+1 | 0.01 ± 0.01 | 0.05 ± 0.00 | 0.15 ± 0.01 |
| | | | M+2 | 0.00 ± 0.00 | 0.07 ± 0.10 | 0.02 ± 0.01 |
| | | | M+3 | 0.00 ± 0.00 | 0.00 ± 0.00 | 0.00 ± 0.00 |
| | | | M+4 | 0.00 ± 0.00 | 0.00 ± 0.00 | 0.00 ± 0.00 |
| | | | M+5 | 0.00 ± 0.00 | 0.00 ± 0.00 | 0.00 ± 0.00 |
| Leu302 | $[M-57]^+$ | C1-C6 | M+0 | 0.98 ± 0.01 | 0.97 ± 0.01 | 0.90 ± 0.02 |
| | | | M+1 | 0.02 ± 0.01 | 0.02 ± 0.01 | 0.09 ± 0.02 |
| | | | M+2 | 0.00 ± 0.00 | 0.01 ± 0.00 | 0.01 ± 0.00 |
| | | | M+3 | 0.00 ± 0.00 | 0.00 ± 0.00 | 0.00 ± 0.00 |
| | | | M+4 | 0.00 ± 0.00 | 0.00 ± 0.00 | 0.00 ± 0.00 |
| | | | M+5 | 0.00 ± 0.00 | 0.00 ± 0.00 | 0.00 ± 0.00 |
| | | | M+6 | 0.00 ± 0.00 | 0.00 ± 0.00 | 0.00 ± 0.00 |
| Leu274 | $[M-85]^+$ | C2-C6 | M+0 | 0.98 ± 0.01 | 0.96 ± 0.01 | 0.90 ± 0.02 |
| | | | M+1 | 0.02 ± 0.01 | 0.03 ± 0.01 | 0.09 ± 0.02 |
| | | | M+2 | 0.00 ± 0.00 | 0.01 ± 0.00 | 0.01 ± 0.01 |
| | | | M+3 | 0.00 ± 0.00 | 0.00 ± 0.00 | 0.00 ± 0.00 |
| | | | M+4 | 0.00 ± 0.00 | 0.00 ± 0.00 | 0.00 ± 0.00 |
| | | | M+5 | 0.00 ± 0.00 | 0.00 ± 0.00 | 0.00 ± 0.00 |
| Lys431 | $[M-57]^+$ | C1-C6 | M+0 | 0.98 ± 0.01 | 0.99 ± 0.02 | 0.91 ± 0.03 |
| | | | M+1 | 0.02 ± 0.01 | 0.01 ± 0.02 | 0.08 ± 0.02 |
| | | | M+2 | 0.00 ± 0.00 | 0.00 ± 0.00 | 0.01 ± 0.01 |
| | | | M+3 | 0.00 ± 0.00 | 0.00 ± 0.00 | 0.00 ± 0.00 |
| | | | M+4 | 0.00 ± 0.00 | 0.00 ± 0.00 | 0.00 ± 0.00 |
| | | | M+5 | 0.00 ± 0.00 | 0.00 ± 0.00 | 0.00 ± 0.00 |
| | | | M+6 | 0.00 ± 0.00 | 0.00 ± 0.00 | 0.00 ± 0.00 |
| Phe336 | $[M-57]^+$ | C1-C9 | M+0 | 0.99 ± 0.00 | 0.97 ± 0.01 | 0.87 ± 0.05 |
| | | | M+1 | 0.00 ± 0.00 | 0.01 ± 0.01 | 0.11 ± 0.03 |
| | | | M+2 | 0.00 ± 0.00 | 0.01 ± 0.00 | 0.02 ± 0.01 |
| | | | M+3 | 0.00 ± 0.00 | 0.01 ± 0.00 | 0.00 ± 0.00 |
| | | | M+4 | 0.00 ± 0.00 | 0.00 ± 0.00 | 0.00 ± 0.00 |
| | | | M+5 | 0.00 ± 0.00 | 0.00 ± 0.00 | 0.00 ± 0.00 |
| | | | M+6 | 0.00 ± 0.00 | 0.00 ± 0.00 | 0.00 ± 0.00 |
| | | | M+7 | 0.00 ± 0.00 | 0.00 ± 0.00 | 0.00 ± 0.00 |

| | | | | | | |
|---|---|---|---|---|---|---|
| | | | M+8 | 0.00 ± 0.00 | 0.00 ± 0.00 | 0.00 ± 0.00 |
| | | | M+9 | 0.00 ± 0.00 | 0.00 ± 0.00 | 0.00 ± 0.00 |
| Phe308 | [M-85]$^+$ | C2-C9 | M+0 | 0.99 ± 0.00 | 0.96 ± 0.01 | 0.88 ± 0.04 |
| | | | M+1 | 0.01 ± 0.00 | 0.02 ± 0.01 | 0.10 ± 0.02 |
| | | | M+2 | 0.00 ± 0.00 | 0.01 ± 0.00 | 0.02 ± 0.01 |
| | | | M+3 | 0.00 ± 0.00 | 0.00 ± 0.00 | 0.00 ± 0.00 |
| | | | M+4 | 0.00 ± 0.00 | 0.00 ± 0.00 | 0.00 ± 0.00 |
| | | | M+5 | 0.00 ± 0.00 | 0.00 ± 0.00 | 0.00 ± 0.00 |
| | | | M+6 | 0.00 ± 0.00 | 0.00 ± 0.00 | 0.00 ± 0.00 |
| | | | M+7 | 0.00 ± 0.00 | 0.00 ± 0.00 | 0.00 ± 0.00 |
| Pro286 | [M-57]$^+$ | C1-C5 | M+0 | 0.00 ± 0.00 | 0.97 ± 0.02 | 0.85 ± 0.02 |
| | | | M+1 | 0.01 ± 0.00 | 0.02 ± 0.01 | 0.13 ± 0.02 |
| | | | M+2 | 0.00 ± 0.00 | 0.01 ± 0.01 | 0.02 ± 0.00 |
| | | | M+3 | 0.00 ± 0.00 | 0.00 ± 0.00 | 0.00 ± 0.00 |
| | | | M+4 | 0.00 ± 0.00 | 0.00 ± 0.00 | 0.00 ± 0.00 |
| | | | M+5 | 0.00 ± 0.00 | 0.00 ± 0.00 | 0.00 ± 0.00 |
| Pro258 | [M-85]$^+$ | C2-C5 | M+0 | 0.99 ± 0.00 | 0.95 ± 0.01 | 0.87 ± 0.02 |
| | | | M+1 | 0.01 ± 0.00 | 0.03 ± 0.01 | 0.12 ± 0.01 |
| | | | M+2 | 0.00 ± 0.00 | 0.01 ± 0.00 | 0.01 ± 0.01 |
| | | | M+3 | 0.00 ± 0.00 | 0.00 ± 0.00 | 0.00 ± 0.00 |
| | | | M+4 | 0.00 ± 0.00 | 0.00 ± 0.00 | 0.00 ± 0.00 |
| Ser390 | [M-57]$^+$ | C1-C3 | M+0 | 0.99 ± 0.01 | 0.97 ± 0.00 | 0.93 ± 0.01 |
| | | | M+1 | 0.01 ± 0.01 | 0.02 ± 0.00 | 0.07 ± 0.01 |
| | | | M+2 | 0.00 ± 0.00 | 0.01 ± 0.00 | 0.00 ± 0.00 |
| | | | M+3 | 0.00 ± 0.00 | 0.00 ± 0.00 | 0.00 ± 0.00 |
| Ser262 | [M-85]$^+$ | C2-C3 | M+0 | 1.00 ± 0.00 | 0.98 ± 0.00 | 0.94 ± 0.01 |
| | | | M+1 | 0.00 ± 0.00 | 0.02 ± 0.00 | 0.06 ± 0.01 |
| | | | M+2 | 0.00 ± 0.00 | 0.00 ± 0.00 | 0.00 ± 0.00 |
| Thr404 | [M-57]$^+$ | C1-C4 | M+0 | 0.99 ± 0.00 | 0.95 ± 0.01 | 0.84 ± 0.01 |
| | | | M+1 | 0.01 ± 0.00 | 0.03 ± 0.01 | 0.15 ± 0.01 |
| | | | M+2 | 0.00 ± 0.00 | 0.01 ± 0.00 | 0.01 ± 0.00 |
| | | | M+3 | 0.00 ± 0.00 | 0.01 ± 0.00 | 0.00 ± 0.00 |
| | | | M+4 | 0.00 ± 0.00 | 0.00 ± 0.00 | 0.00 ± 0.00 |
| Thr376 | [M-85]$^+$ | C2-C4 | M+0 | 1.00 ± 0.00 | 0.96 ± 0.01 | 0.86 ± 0.01 |
| | | | M+1 | 0.00 ± 0.00 | 0.03 ± 0.01 | 0.13 ± 0.01 |
| | | | M+2 | 0.00 ± 0.00 | 0.01 ± 0.00 | 0.01 ± 0.00 |
| | | | M+3 | 0.00 ± 0.00 | 0.00 ± 0.00 | 0.00 ± 0.00 |
| Tyr466 | [M-57]$^+$ | C1-C9 | M+0 | 0.99 ± 0.01 | 0.96 ± 0.02 | 0.84 ± 0.04 |
| | | | M+1 | 0.01 ± 0.01 | 0.02 ± 0.01 | 0.13 ± 0.02 |
| | | | M+2 | 0.00 ± 0.00 | 0.00 ± 0.01 | 0.02 ± 0.01 |
| | | | M+3 | 0.00 ± 0.01 | 0.02 ± 0.00 | 0.00 ± 0.01 |
| | | | M+4 | 0.00 ± 0.00 | 0.00 ± 0.00 | 0.00 ± 0.00 |
| | | | M+5 | 0.00 ± 0.00 | 0.00 ± 0.00 | 0.00 ± 0.00 |
| | | | M+6 | 0.00 ± 0.00 | 0.00 ± 0.00 | 0.00 ± 0.00 |

| | | | | | | |
|---|---|---|---|---|---|---|
| | | | M+7 | 0.00 ± 0.00 | 0.00 ± 0.00 | 0.00 ± 0.00 |
| | | | M+8 | 0.00 ± 0.00 | 0.00 ± 0.00 | 0.00 ± 0.00 |
| | | | M+9 | 0.00 ± 0.00 | 0.00 ± 0.00 | 0.00 ± 0.00 |
| Tyr438 | [M-85]+ | C2-C9 | M+0 | 0.88 ± 0.05 | 0.86 ± 0.05 | 0.73 ± 0.07 |
| | | | M+1 | 0.00 ± 0.01 | 0.02 ± 0.01 | 0.11 ± 0.04 |
| | | | M+2 | 0.00 ± 0.00 | 0.01 ± 0.01 | 0.00 ± 0.00 |
| | | | M+3 | 0.00 ± 0.00 | 0.00 ± 0.00 | 0.00 ± 0.00 |
| | | | M+4 | 0.00 ± 0.00 | 0.00 ± 0.00 | 0.00 ± 0.00 |
| | | | M+5 | 0.00 ± 0.00 | 0.00 ± 0.00 | 0.00 ± 0.00 |
| | | | M+6 | 0.00 ± 0.00 | 0.00 ± 0.00 | 0.00 ± 0.00 |
| | | | M+7 | 0.00 ± 0.00 | 0.00 ± 0.00 | 0.00 ± 0.00 |
| | | | M+8 | 0.12 ± 0.06 | 0.11 ± 0.05 | 0.16 ± 0.10 |
| Val288 | [M-57]+ | C1-C5 | M+0 | 0.99 ± 0.01 | 0.96 ± 0.00 | 0.87 ± 0.02 |
| | | | M+1 | 0.01 ± 0.01 | 0.02 ± 0.00 | 0.11 ± 0.02 |
| | | | M+2 | 0.00 ± 0.00 | 0.01 ± 0.00 | 0.01 ± 0.00 |
| | | | M+3 | 0.00 ± 0.00 | 0.01 ± 0.00 | 0.00 ± 0.00 |
| | | | M+4 | 0.00 ± 0.00 | 0.00 ± 0.00 | 0.00 ± 0.00 |
| | | | M+5 | 0.00 ± 0.00 | 0.00 ± 0.00 | 0.00 ± 0.00 |
| Val260 | [M-85]+ | C2-C5 | M+0 | 0.98 ± 0.01 | 0.94 ± 0.01 | 0.86 ± 0.02 |
| | | | M+1 | 0.02 ± 0.01 | 0.04 ± 0.01 | 0.13 ± 0.02 |
| | | | M+2 | 0.00 ± 0.00 | 0.02 ± 0.01 | 0.01 ± 0.00 |
| | | | M+3 | 0.00 ± 0.00 | 0.00 ± 0.00 | 0.00 ± 0.00 |
| | | | M+4 | 0.00 ± 0.00 | 0.00 ± 0.00 | 0.00 ± 0.00 |

***Table S13** The mass isotopomer distributions (MIDs) [represented as mean ± standard deviation (SD); n = 4] in valid amino acid fragments ([M-57]+ and [M-85]+) of Alternaria burnsii grown in Potato infusion broth supplemented with 99% [$^{12}$C]glucose, 20% [U-$^{13}C_6$]glucose and 99 % [1-$^{13}$C]glucose [Ala: Alanine; Asp: Aspartic acid; Glu: Glutamic acid; Gly: Glycine; His: Histidine; Ile: Isoleucine; Leu: Leucine; Lys: Lysine; Phe: Phenylalanine; Pro: Proline; Ser: Serine; Thr: Threonine; Tyr: Tyrosine; Val: Valine].*

### SUPPLEMENTARY REFERENCES


1. Lu, Y., Ye, C., Che, J., Xu, X., Shao, D., Jiang, C., Liu, Y. & Shi, J.: Genomic sequencing, genome-scale metabolic network reconstruction, and in silico flux analysis of the grape endophytic fungus *Alternaria sp*. MG1. Microb Cell Fact, 18, 1–16 (2019).
2. Mohinudeen, I. A. H. K., Kanumuri, R., Soujanya, K. N., Shaanker, R. U., Rayala, S. K. & Srivastava, S.: Sustainable production of camptothecin from an *Alternaria sp*. isolated from *Nothapodytes nimmoniana*. Scientific Reports 2021 11:1, 11, 1–11 (2021).
